# Endothelial calcium firing mediates extravasation of metastatic tumor cells

**DOI:** 10.1101/2024.03.28.587188

**Authors:** Marina Peralta, Amandine Dupas, Annabel Larnicol, Olivier Lefebvre, Ruchi Goswami, Tristan Stemmelen, Anne Molitor, Raphael Carapito, Salvatore Girardo, Naël Osmani, Jacky G. Goetz

**Author notes:** Materials and correspondence, lead author: Jacky G. Goetz, Naël Osmani, Marina Peralta, Web: www.goetzlab.fr.

## Abstract

Metastatic dissemination is driven by genetical, biochemical and biophysical cues that favor the distant colonization of organs and the formation of life-threatening secondary tumors. We have previously demonstrated that endothelial cells (ECs) actively remodel during extravasation by enwrapping arrested tumor cells (TCs) and extrude them from the vascular lumen while maintaining perfusion. In this work, we dissect the cellular and molecular mechanisms driving endothelial remodeling. Using high-resolution intravital imaging in zebrafish embryos, we demonstrate that the actomyosin network of ECs controls tissue remodeling and subsequent TC extravasation. Furthermore, we uncovered that this cytoskeletal remodeling is driven by altered endothelial- calcium (Ca^2+^) signaling caused by arrested TCs. Accordingly, we demonstrated that inhibition of voltage-dependent calcium L-type channels impairs extravasation. Lastly, we identified P2X4, TRP, and Piezo1 mechano-gated Ca^2+^ channels as key mediators of the process. These results further highlight the central role of endothelial remodeling during extravasation of TCs and open avenues for successful therapeutic targeting.

**Graphical Abstract:** 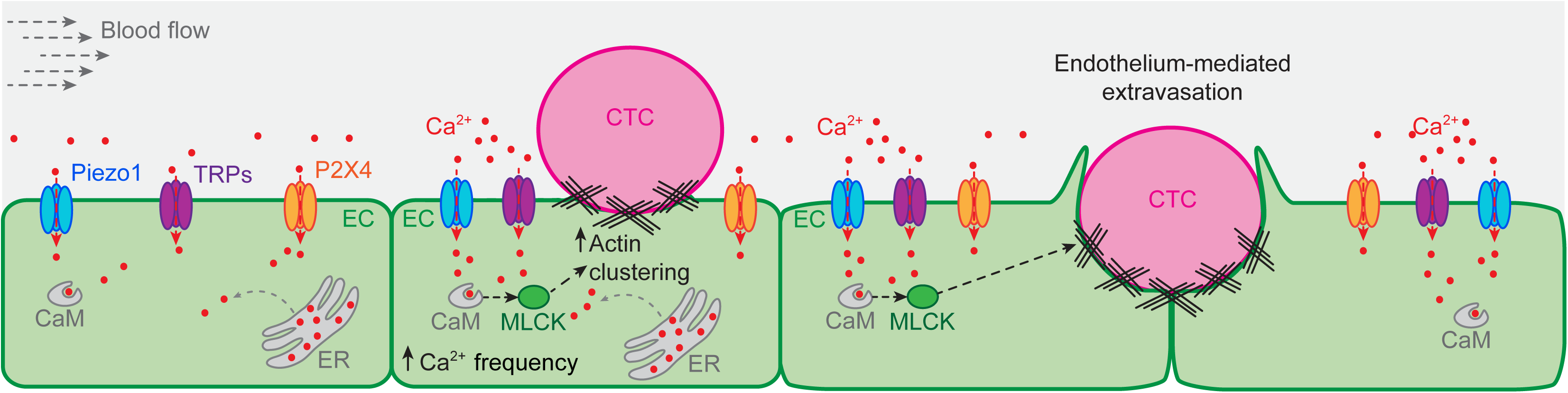

## INTRODUCTION

Metastatic colonization of distant organs occurs during tumorigenic progression and is the main reason for cancer-related death^1–3^. Metastatic dissemination relies on biochemical and biophysical cues either from TCs or from their direct microenvironment that will dictate organotropism and the outcome of distant organ colonization. The adhesion molecule repertoire^4,5^, the secretome^6–9^ of TCs, and the vascular features^10^, together with biomechanical characteristics from the disseminating cells and the microenvironment^11–13^ impact the early steps of TC dissemination including extravasation and early seeding of cancer cells in distant organs.

The vasculature is central for the dissemination of circulating tumor cells (CTCs) not only as a passive transport system but also as an active player in the colonization of distant sites^14^. The perivascular niche is highly colonized by metastatic cells becoming a central hub instructing metastatic outcome in most organs^15–18^. The vasculature also dictates the arrest of CTCs either by direct occlusion^19,20^ or by defining hemodynamic patterns that are compatible with CTC adhesion to the luminal endothelium^5,21^. As a barrier between the blood stream and the organs, the endothelium must be crossed by CTCs and thus imposes their extravasation efficiency. The mechanical features of endothelial cells (ECs) and their adhesion repertoire are elemental in fine-tuning extravasation efficiency^4,22^. In addition to diapedesis, we and others have described alternative routes to exit the vasculature. Specifically, the active remodeling of the endothelium in response to CTC arrest promotes indiscriminately the extravasation of single CTCs or highly metastatic clusters notably during brain metastasis^21,23–25^. This endothelial indoctrination requires not only the biomechanical input of flow forces activating VEGFR signaling but also secreted factors from arrested CTCs such as MMP9^21,24,25^. This suggests that biochemical and biophysical cues exerted on EC luminal side need to be transduced to mediate intraluminal vascular remodeling and thus facilitate CTC successful extravasation. Calcium (Ca^2+^) is an important second messenger that integrates external biochemical and biophysical signals and is essential in angiogenesis and vascular homeostasis and function^26–28^. Ca^2+^ is known to drive cell shape changes by remodeling the actomyosin cytoskeleton. More specifically, Ca^2+^ influx has been shown to promote actomyosin contraction and remodeling through several mechanism, including the activation of myosin light chain kinase^29,30^. We thus hypothesized that Ca^2+^ signaling is central to CTC extravasation during metastatic progression, particularly by favoring endothelial cytoskeletal actomyosin reorganization. In the present work, we used high-resolution intravital imaging in a zebrafish experimental metastasis model to highlight the role of actomyosin cytoskeleton reorganization in endothelial remodeling. Taking advantage of a genetically encoded endothelial-Ca^2+^ influx reporter zebrafish line, we observed that upon CTC arrest and throughout the extravasation process, ECs displayed increased Ca^2+^ firing, which promoted actomyosin rearrangements essential for intravascular endothelial remodeling. Furthermore, we demonstrated that Ca^2+^ is required for endothelial indoctrination by TCs and thus their extravasation. Lastly, we identified P2X4, TRP, and Piezo1 mechano-gated channels as mediators of the process.

## RESULTS

### TCs stimulate endothelial actomyosin-dependent remodeling to extravasate

We and others previously described the active participation of the endothelium during CTC extravasation in a process that we named intraluminal endothelial remodeling. Such process occurs for extravasation of both single and cluster CTCs and secures efficient metastatic seeding^21,23,25^. However, the cellular and molecular mechanisms at play during such morphological remodeling of the endothelium remain unclear. Here, we aimed to improve the cellular and temporal resolution to uncover the complex cytoskeletal changes occurring within ECs, while providing an additional dissection of its molecular control. We exploited single cell resolution intravital live imaging and relied on our metastasis experimental model in the zebrafish embryo^31^. Using the *Tg(fli:lifeAct-eGFP;flk:nls-mCherry)* reporter line, which allows single endothelial cell resolution with EC nuclei labelled (mCherry), we assessed cytoskeletal actomyosin dynamics at single cell level with a polymerized actin probe (LifeAct-eGFP). We injected mouse mammary carcinoma D2A1 cells in the duct of Cuvier of 2 days post fertilization (dpf) zebrafish embryos and performed time lapse imaging every 45 min during 11 h (Fig.1A). When probing CTC arrest up to its extravasation, we noticed that it was accompanied by a massive EC actin clustering (Fig.1B-C; Movie 1). As the embryonic vasculature naturally develops, the EC cytoskeletal activity is highly dynamic and actin clusters are visible through time. In order to provide a meaningful quantification of TC-based Actin clustering, we designed a quantitative analysis pipeline that accounted for the intrinsic EC cytoskeletal reorganization. To that end, we calculated the cumulative measurements of the EC-actin clusters area over time in regions of interest (ROIs) around CTCs and ROIs in equivalent EC regions lacking CTCs. We then normalized each CTC-ROI measurement to its no- CTC-ROI control. By doing so, we can safely distinguish the specific effect of CTCs on the endothelial actomyosin cytoskeleton (Fig.1B’). This quantitative analysis demonstrated that CTCs induce additional EC actin clustering independently of the actomyosin cytoskeletal dynamics of the embryonic endothelium (Fig.1D). Importantly, EC actin clustering during CTC arrest and extravasation was observed not only in small-sized blood vessels such as inter-somitic vessels (ISVs, Fig.1C), but also in bigger ones such as the caudal vein (CV) and the dorsal aorta (DA, Fig.S1A). Furthermore, to test the analysis pipeline robustness, we drew two no-CTC-ROIs for each time point in regions where CTCs usually arrested and assigned them randomly to group A or B. Then, we performed the previously described analysis normalizing group A to B and conversely group B to A. If the differences that we observed between CTC- and no-CTC-ROIs were unspecific, they would appear between these randomized no-CTC-ROIs. However, this analysis led to non-significant differences on EC actin clustering in the absence of arrested TCs, further stressing the high specificity of our analysis pipeline (Fig.S1B). This innovative area-based analysis provided the first quantitative description of the EC actomyosin cytoskeletal dynamics upon CTC arrest and up to their extravasation.

**Figure 1:**
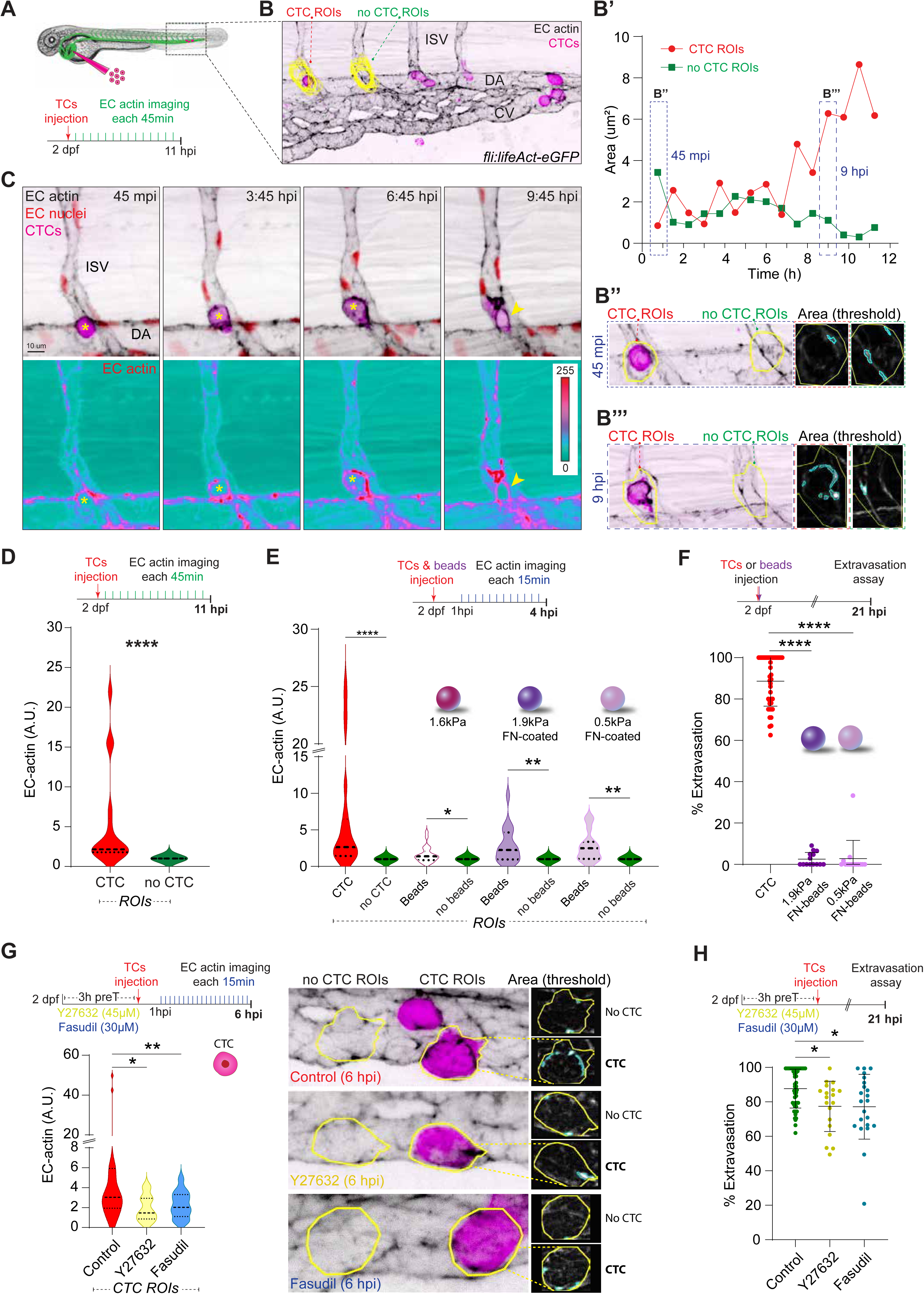
TCs stimulate endothelial actomyosin-dependent remodeling to extravasate. **A.** Schematic representation of the experimental design. **B.** Example of EC-actin quantification: Yellow CTC- and no-CTC- ROIs are projected over the first time point of the time-lapse. Graph (**B’**) shows the EC-actin area through time. Example of the analysis pipeline selections for CTC- and no-CTC- ROIs (yellow line) at 45 mpi (**B’’**) and 9 hpi (**B’’’**). **C.** Time-lapse z-stack projections show a CTC extravasating from an ISV. Bellow, individual actin channel is displayed using ice LUT to facilitate the visualization of signal intensity, where green is the minimum and red is the maximum. For all the images, yellow asterisks label intravascular CTC, yellow arrowhead mark the extravasated CTC. **D.** Schematic representation of the experimental design. Graph shows the EC actin clustering for each pair of ROIs (CTC and no-CTC) throughout the time lapse in arbitrary units (A.U., Wilcoxon matched-pairs signed rank test, p-value <0.0001, 3 embryos, 17 CTC- and no-CTC- ROIs measured for 11 hpi time-lapse). **E.** Schematic representation of the experimental design. Graph shows the EC actin clustering for each pair of ROIs (CTC and no-CTC or beads and no-beads, respectively) throughout the time lapse in arbitrary units (A.U., Wilcoxon matched-pairs signed rank test, CTCs: p-value <0.0001; 1.6kPa beads: p-value 0.019; 1.9kPa fibronectin-coated beads: p-value 0.0028; 0.5kPa fibronectin-coated beads: p-value 0.0034, CTCs: 3 embryos, 17 CTC- and no-CTC- ROIs measured; 1.6kPa beads: 8 embryos, 23 bead- and no-bead- ROIs measured; 1.9kPa fibronectin-coated beads: 5 embryos, 18 bead- and no-bead- ROIs measured; 0.5kPa fibronectin-coated beads: 8 embryos, 15 bead- and no-bead- ROIs measured for 4 hpi time-lapse). **F.** Schematic representation of the experimental design. Graph shows CTCs, 1.9kPa- and 0.5kPa- fibronectin-coated beads percentage of extravasation at 21 hpi (Kruskal-Wallis test multiple comparison p values < 0.0001; CTCs: 42 embryos; 1.9kPa: 14 embryos; 0.5kPa: 14 embryos). **G.** Schematic representation of the experimental design. Graph shows the EC actin clustering for control, Y27632- or Fasudil-treated embryos throughout the time lapse in arbitrary units (A.U., Kruskal-Wallis test multiple comparison control vs. Y27632 p value 0.048; control vs. Fasudil p value 0.0028; control: 6 embryos, 28 CTC-ROIs measured; Y27632: 4 embryos, 13 CTC-ROIs measured; Fasudil: 8 embryos, 13 CTC-ROIs measured for 6 hpi time-lapse). Images show representative examples of the analysis pipeline selections for CTC- and no-CTC- ROIs (yellow line) at 6 hpi for each condition. **H.** Schematic representation of the experimental design. Graph shows control, Y27632- or Fasudil-treated embryos percentage of extravasation (Kruskal-Wallis test multiple comparison control vs. Y27632 p value 0.024; control vs. Fasudil p value 0.031; control: 48 embryos; Y27632: 19 embryos; Fasudil: 22 embryos).

Upon arrest, a TC exerts mechanical and chemical stimuli on ECs^12,13^. Importantly, mechanical cues are known mediators of actin remodelling^32^, including intravascular pressure, which had been shown to induce actin polymerization in vascular smooth muscle^33^. To explore the extent to which a mechanical stimulus exerted on ECs (i.e. locally affecting the EC plasma membrane and cytoskeleton, and/or altering the local flow profile) is sufficient to induce EC actomyosin cytoskeletal remodeling, we leveraged three types of polyacrylamide microgel beads to mimic the physical cues of an arrested TC^34^. Intravascular TCs display a rounded shape, ∼15 um in diameter, and 0.75-0.85 kPa^21,35^. All the beads displayed similar diameter, but different stiffness (stiffer or softer) profiles than of TCs of choice (D2A1 cells). We compared the effect of beads and TCs on EC actin remodeling by performing time lapse imaging every 15 min during 4 h. Inert polyacrylamide beads (∼15.2 μm diameter and 1.6 kPa) elicited a very low EC actin clustering response (Fig.1E; Fig.S1C and D). While this result highlights the robustness and sensitivity of the analysis pipeline, it mostly demonstrates that mechanical stimuli exerted on ECs by arrested objects without a molecular engagement only produced a weak cytoskeletal response. In order to model weak adhesions between beads and the endothelium, we relied on fibronectin (FN)-coated beads that would either engage integrin α5β1 on the endothelial luminal side^36,37^ or to fibronectin deposits at the luminal surface of ECs as we previously reported^5^. We tested two types of FN-coated beads displaying ∼16.5 μm diameter and 1.9 kPa (stiffer than D2A1) or 0.5 kPa (softer than D2A1). The stiffness range aimed at exploring the possible role of CTC mechanical properties. However, the range of EC actin clustering values upon bead-EC contact were lower than the CTC-EC-driven (Fig.1E). These results demonstrate that the EC actomyosin reorganization specifically results from combined mechanical and biochemical stimuli exerted by arrested TCs.

Aiming to further demonstrate that EC actomyosin rearrangements were directly correlated and involved in extravasation, we tested the extravasation capability of both FN-coated bead types, as they elicited a stronger EC cytoskeletal response than non-coated beads. At 21 hpi, most FN-coated beads of any stiffness failed to extravasate: 1.9 kPa-beads: 0.74% extravasation (3 extravasated beads of 405 injected in 14 embryos); 0.5 kPa-beads: 4.3% extravasation (17 extravasated beads of 398 injected in 14 embryos, Fig.1F). Taken together, these results identified EC cytoskeletal reorganization as an important driver for endothelial remodeling during TC extravasation and exposed the need for strong molecular engagement and biochemical cues provided by CTCs in order to indoctrinate ECs.

As minor EC actin clustering is insufficient to drive extravasation of inert beads, we next assessed the extent to which EC cytoskeletal reorganization is needed for CTC extravasation and therefore, in metastasis formation. To do so, we targeted Rho-associated kinase (ROCK), which is known to play a major role in mediating actomyosin cytoskeleton rearrangements^38,39^, with clinically-relevant pharmacological approaches. We disrupted actin dynamics using the ROCK1 and 2 inhibitor Y-27632^40^ as well as fasudil, a ROCK inhibitor and vasodilator (calcium antagonist) approved for clinical purposes in Japan and China^41,42^. When probing EC actin dynamics in control and treated-embryos with CTCs, we observed that the latter triggered significantly weaker EC cytoskeletal response (Fig.1G). We further assessed extravasation upon actomyosin contraction inhibition and observed decreased CTC extravasation in Y-27632- and fasudil-treated embryos (Fig.1H). Altogether, this demonstrates that TC rapidly engage with ECs and stimulate actomyosin contraction that sets extravasation-prone morphological changes.

### Arrest of CTCs triggers endothelial calcium firing

We next aimed to understand the molecular pathways regulating EC actomyosin rearrangements upon CTC arrest. Calcium is an important cell secondary messenger that regulates cell shape and motility by controlling cytoskeletal actomyosin in response to mechanical stimuli^26,43,44^. We thus hypothesized that intravascular endothelial remodeling could be initiated by TC-dependent Ca^2+^ firing. To assess this hypothesis, we took advantage of the *Tg(fli:GalFF; UAS:GCAMP7a)*, a reporter line to study, for the first time in the context of metastatic extravasation, the dynamics of intracellular Ca^2+^ in ECs *in vivo*^45^. GCaMP7a (GFP-based Ca^2+^ probe) is an engineered GFP that increases fluorescence upon Ca^2+^ influx^46^. We performed high-speed imaging (for 5 min) on injected and non-injected (henceforth controls) embryos at 2 hpi (Fig.2A). Such timing (2hpi) allows us to probe recently arrested CTC, where the contact can still be disengaged and while some TCs are still circulating^5^. We characterized endothelial Ca^2+^ oscillations according to frequency (*i.e.*, number of Ca^2+^ oscillations per EC) and amplitude (i.e., maximum intensity of Ca^2+^ oscillations for each EC) in the DA and CV in control and injected embryos (Fig. 2B-D; Fig.S2A and B). Strikingly, arrest and engagement of TCs with ECs trigger endothelial Ca^2+^ firing, where frequency, but not amplitude, was significantly increased (Fig.S2B; Fig.2B-D; Movie 2 and 3). When distinguishing ECs that are in direct contact with arrested TCs from ECs that are not in direct contact with arrested TCs, we observed an increased endothelial Ca^2+^ firing in close proximity to arrested TCs (Fig.S2C). Although such observation did not reach statistical significance, it suggests that TCs can indoctrinate endothelial cell signaling locally, either by direct interaction or by short-range paracrine signaling. When probing endothelial Ca^2+^ firing at 6 hpi, when most CTCs are arrested and extravasation-prone intraluminal endothelial remodeling is engaged^24^, we again observed increased endothelial Ca^2+^ firing, with no effect on the amplitude of the oscillations and independently of their proximity to the arrested TCs (Fig.S2D). As endothelial Ca^2+^ firing could be, in theory, solely triggered by mechanical inputs, we explored such scenario and probed Ca^2+^ dynamics with the three types of inert polyacrylamide microgel beads previously described. Interestingly, endothelial Ca^2+^ remained unperturbed independently of the stiffness (0.5, 1.6 or 1.9 kPa) or the coating (FN) profile of the beads (Fig.2E-J; Fig.S2E). Taken together, these results suggest that the arrest and engagement of CTCs are sufficient to initiate local endothelial Ca^2+^ firing independently of cell mechanics.

**Figure 2:**
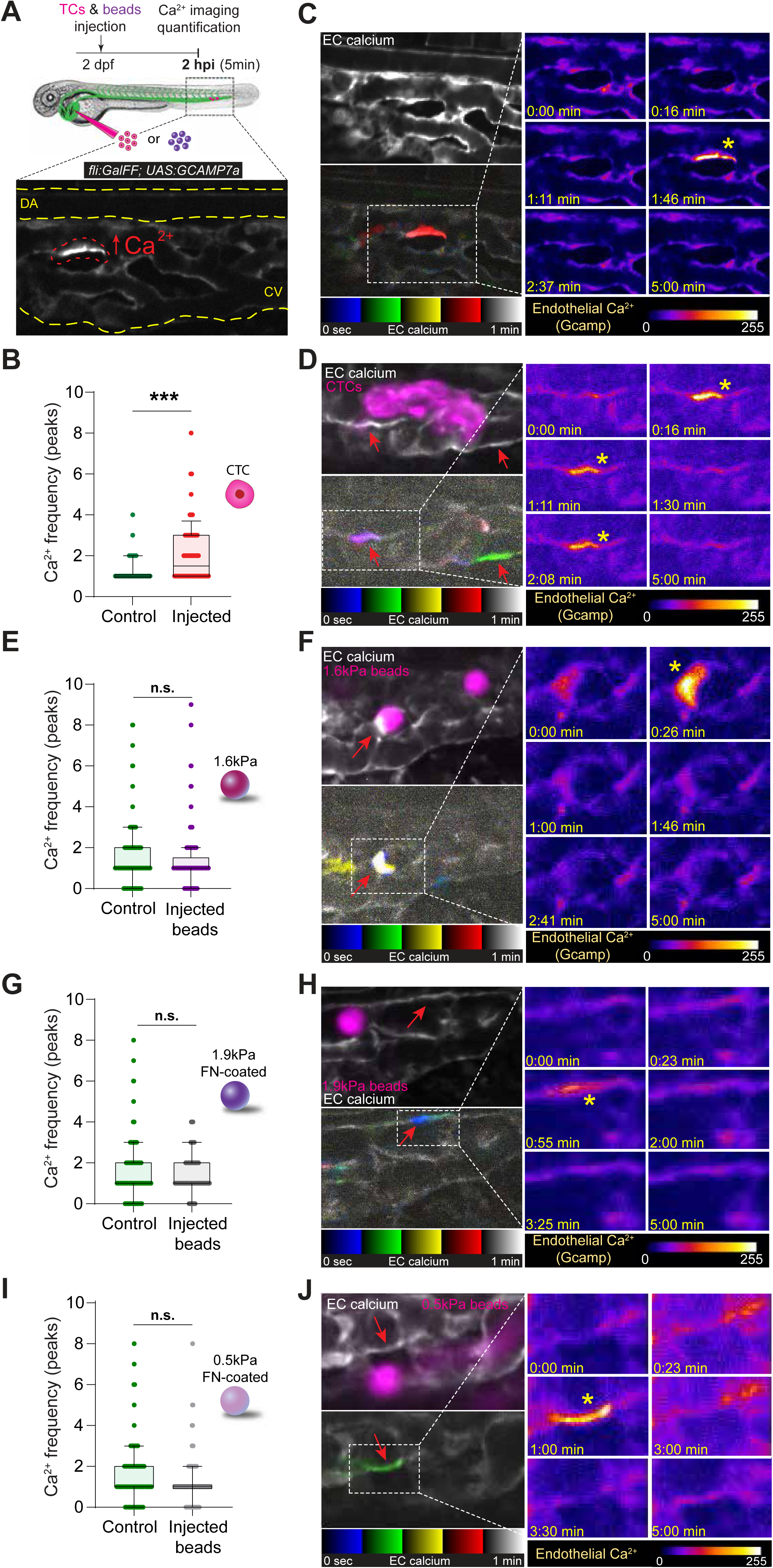
Arrest of CTCs triggers endothelial calcium firing. **A.** Schematic representation of the experimental design. Image illustrates calcium firing in an EC. **B.** Graph shows EC calcium firing frequency (number of peaks) in non-injected (control) and injected embryos at 2 hpi (Mann Whitney test, p-value 0.0002; controls: 5 embryos, 52 cells quantified; injected: 5 embryos, 82 cells quantified). **C.** Confocal section showing EC calcium signaling (white). Bellow, EC calcium imaging displayed using temporal color code for one minute in a control embryo showing one EC calcium firing (red). Right panel: zooms of representative timepoints extracted from 5 min continuous confocal imaging. Signal displayed using fire LUT to facilitate the visualization of signal intensity, where blue is the minimum and white is the maximum. Asterisk highlights an EC calcium firing. **D.** Confocal section showing EC calcium signaling (white) and CTCs (magenta). Bellow, EC calcium imaging displayed using temporal color code for one minute in a CTCs-injected embryo showing two ECs calcium firings (one EC showing one calcium firing in green; another EC in purple because calcium fired twice, one in the blue and one in the red display). Right panel: zooms of representative timepoints extracted from 5 min continuous imaging. Signal displayed using fire LUT to facilitate the visualization of signal intensity, where blue is the minimum and white is the maximum. Asterisks highlight EC calcium firings. **E.** Schematic representation of the experimental design. Graph shows 1.6kPa beads injections (Mann Whitney test, p value 0.38; controls: 7 embryos, 184 cells quantified; injected: 8 embryos, 153 cells quantified). **F.** Confocal section showing EC calcium signaling (white) and 1.6kPa beads (magenta). Bellow, EC calcium imaging displayed using temporal color code for one minute in an injected embryo showing one EC calcium firings (white). Right panel: zooms of representative timepoints extracted from 5 min continuous imaging. Signal displayed using fire LUT to facilitate the visualization of signal intensity, where blue is the minimum and white is the maximum. Asterisk highlights an EC calcium firing. **G.** Graph shows 1.9kPa-fibronectin-coated beads injections (Mann Whitney test, p-value 0.19; controls: 8 embryos, 177 cells quantified; injected: 7 embryos, 182 cells quantified). **H.** Confocal section showing EC calcium signaling (white) and 1.9kPa – FN-coated beads (magenta). Bellow, EC calcium imaging displayed using temporal color code for one minute in an injected embryo showing one EC calcium firing (blue). Right panel: zooms of representative timepoints extracted from 5 min continuous imaging. Signal displayed using fire LUT to facilitate the visualization of signal intensity, where blue is the minimum and white is the maximum. Asterisk highlights an EC calcium firing. **I.** Graph shows 0.5kPa-fibronectin-coated beads injections (Mann Whitney test, p value 0.08; controls: 4 embryos, 99 cells quantified; injected: 5 embryos, 113 cells quantified). **J.** Confocal section showing EC calcium signaling (white) and 0.5kPa – FN-coated beads (magenta). Bellow, EC calcium imaging displayed using temporal color code for one minute in an injected embryo showing one EC calcium firing (green). Right panel: zooms of representative timepoints extracted from 5 min continuous imaging. Signal displayed using fire LUT to facilitate the visualization of signal intensity, where blue is the minimum and white is the maximum. Asterisk highlights an EC calcium firing. In all images, red arrows point at selected ECs. All graphs are boxplots (upper/lower quartile, median, bars show the 10%-90% range), each dot represents an individual EC.

### Impairing endothelial Ca^2+^ signaling alters metastatic extravasation

Prompted by our observations that Ca^2+^ firing and actomyosin contractility sets intravascular remodeling upon CTC arrest (2 hpi) and at early stages of extravasation (6 hpi), we next aimed at testing whether targeting endothelial Ca^2+^ could impair extravasation of TCs. To that end, we first took advantage of nifedipine, an FDA-approved calcium antagonist that inhibits calcium influx by blocking voltage-dependent L-type channels^47^. When probing TC-dependent actin dynamics in *Tg(fli:lifeActin-eGFP;flk:nls-mCherry)* embryos pre-treated with Nifedipine for 1 h, we observed that Ca^2+^inhibition (both frequency and amplitude were perturbed (Fig.S3A) was sufficient to impair endothelial cytoskeletal response (Fig.3A). Furthermore, and as expected, such effect had a profound impact on the extravasation of TCs whose efficiency was reduced 4-fold in embryos treated with nifedipine, demonstrating that TC-mediated Ca^2+^ firings trigger cytoskeletal-based intraluminal remodeling that shapes metastatic extravasation (Fig.3B). As blood flow additionally tunes arrest and extravasation of CTCs by directly impacting intraluminal remodeling^21^ and is likely to impact endothelial Ca^2+^ firing, we set out to discriminate such contributions building on the clinical usage of nifedipine, an anti- hypertensive treatment decreasing the cardiac pacemaker activity. We first compared the extravasation efficiency (21hpi) of TCs in embryos subjected either to nifedipine- or lidocaine- treatments (as previously described^21^). While both treatments significantly reduce pacemaker activity (Nifedipine: ∼120 beats per minute (bpm); Lidocaine: ∼117 bpm; control: ∼180 bpm) and thus blood flow, nifedipine was more potent in reducing extravasation and actin dynamics (Fig.S3B-D) suggesting that direct targeting of endothelial Ca^2+^ signaling has a strong potential in impairing extravasation. This is further demonstrated in experiments where we probed Ca^2+^ dynamics of lidocaine-treated embryos and observed that, although reducing blood flow has a direct effect on TC extravasation, it has no impact on endothelial Ca^2+^ firing (Fig.S3A).

**Figure 3:**
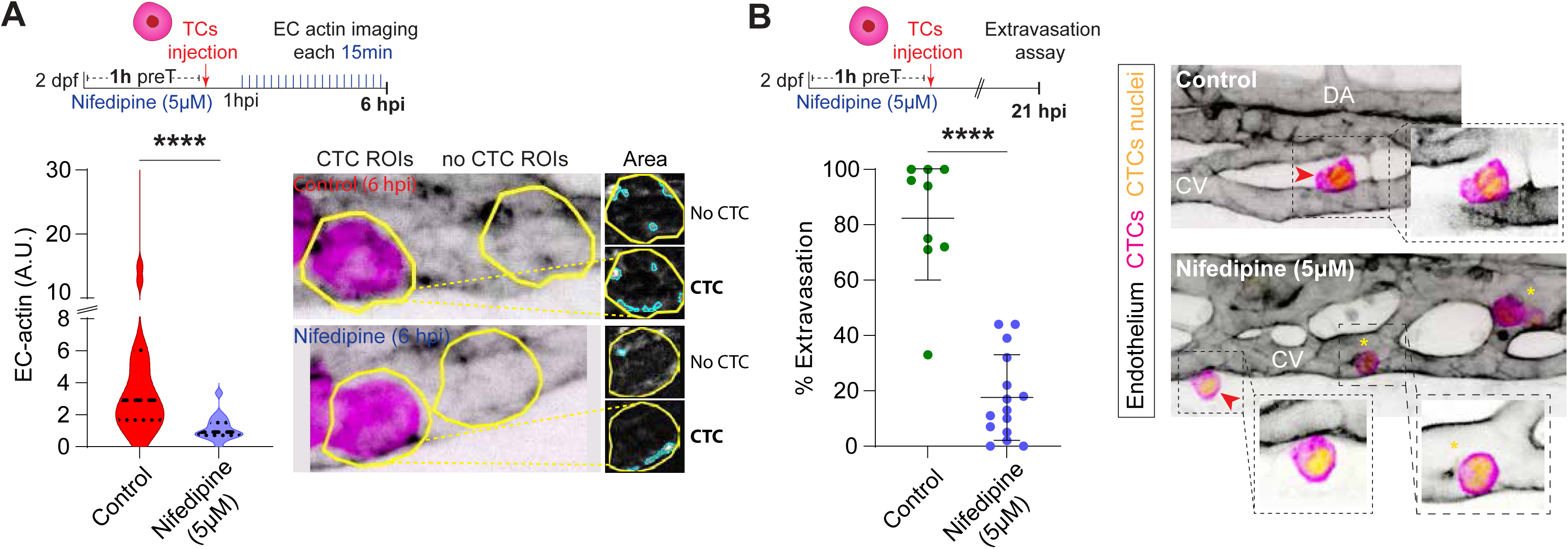
Impairing Ca^2+^ signaling prevents metastatic extravasation. **A.** Schematic representation of the experimental design. Graph shows EC actin clustering for control (CTC ROIs from non-treated embryos) and nifedipine-treated embryos throughout the time lapse in arbitrary units (A.U., Mann Whitney test, p-value 0.0001; control: 3 embryos, 19 CTC- ROIs measured; nifedipine: 2 embryos, 18 CTC- ROIs measured for 6 hpi time-lapse). Images show representative examples of the analysis pipeline selections for CTC- and no-CTC- ROIs (yellow line) at 6 hpi for each condition. **B.** Schematic representation of the experimental design. Graph shows the percentage of CTC extravasation in control and nifedipine-treated embryos at 21 hpi (Mann Whitney test, p-value 0.0001; control: 9 embryos; nifedipine-treated: 15 embryos). Confocal z-stack projections displaying representative examples of control and nifedipine-treated embryos. EC channel is displayed using inverted LUT to facilitate visualization. Zoom boxes show a single confocal plane to improve visualization of intravascular (yellow asterisks) and extravascular (red arrowhead) CTCs.

To further demonstrate that nifedipine effect is not limited to the D2A1 mouse cell line, we also tested its ability to impair the extravasation of a patient-derived human metastatic melanoma cell line (WM983B). We observed that, while milder, nifedipine treatment was still effective at decreasing extravasation, suggesting that its effect does not depend on cancer type (Fig.S3E).

### Metastatic extravasation is impaired by targeting endothelial mechano-gated calcium channels

Encouraged by these results suggesting that endothelial Ca^2+^ signaling directly controls metastatic extravasation, we aimed to provide a molecular identification of the channels responsible for such effect. ECs contain stretch- and shear - activated Ca^2+^ channels that rapidly react to mechanical stimuli applied locally leading to Ca^2+^ influx^48^. These mechano-gated receptors include purinergic P2X receptors, transient receptor potential (TRP), and Piezo channels^27^. ATP released by ECs in response to flow was shown to drive Ca^2+^ influx through P2X4 *in vitro*^49,50^. In the zebrafish, mechanical forces activate EC purinergic channels P2X1, P2X4 and P2X7 modulating Ca^2+^ influx in response to changes in extracellular ATP levels during cardiac valve formation^51^. ECs express a variety of TRP channels, which play an important role in different endothelial functions, including vascular permeability and sensing hemodynamic and chemical changes^52^. Likewise, Piezo1 channels in ECs respond to blood flow by shear-stress-evoked Ca^2+^ influx, which determine the vascular development and architecture in mouse^53,54^. In addition, EC Piezo1 is required for flow-induced ATP release and subsequent purinergic receptors activation that controls blood pressure^55^. We thus leveraged 5-BDBD, a selective P2X4 receptor antagonist; A-438079, a selective P2X7 receptor antagonist; GsMTx4, a spider venom peptide that selectively inhibits cationic mechanosensitive channels belonging to TRP and Piezo families; and Dooku1, a Piezo1 antagonist in ECs^51,56,57^ and subjected our experimental metastatic extravasation read-out (21 hpi) to these treatments. Interestingly, while 5-BDBD, GsMTx4, and Dooku1 significantly reduced metastatic extravasation (Controls = ∼88%, 5-BDBD = ∼68%; GsMTx4 = ∼68%; Dooku1 = ∼72%, Fig. 4A-D), A-438079 (∼84%) had no effect (Fig.S3F). Moreover, we also observed a strong impairment of CTC-induced endothelial actin clustering in embryos treated with altered 5-BDBD, GsMTx4 or Dooku1 (Fig. 4E-H). This confirms a direct role of Ca^2+^ mechanosignaling in CTC-induced actin clustering and subsequent endothelial driven extravasation. Taken together, these results suggest that metastatic extravasation can be controlled by targeting endothelial calcium channels such as TRP and/or Piezo, together with P2X4, providing promising targets for anti-metastasis treatments.

**Figure 4:**
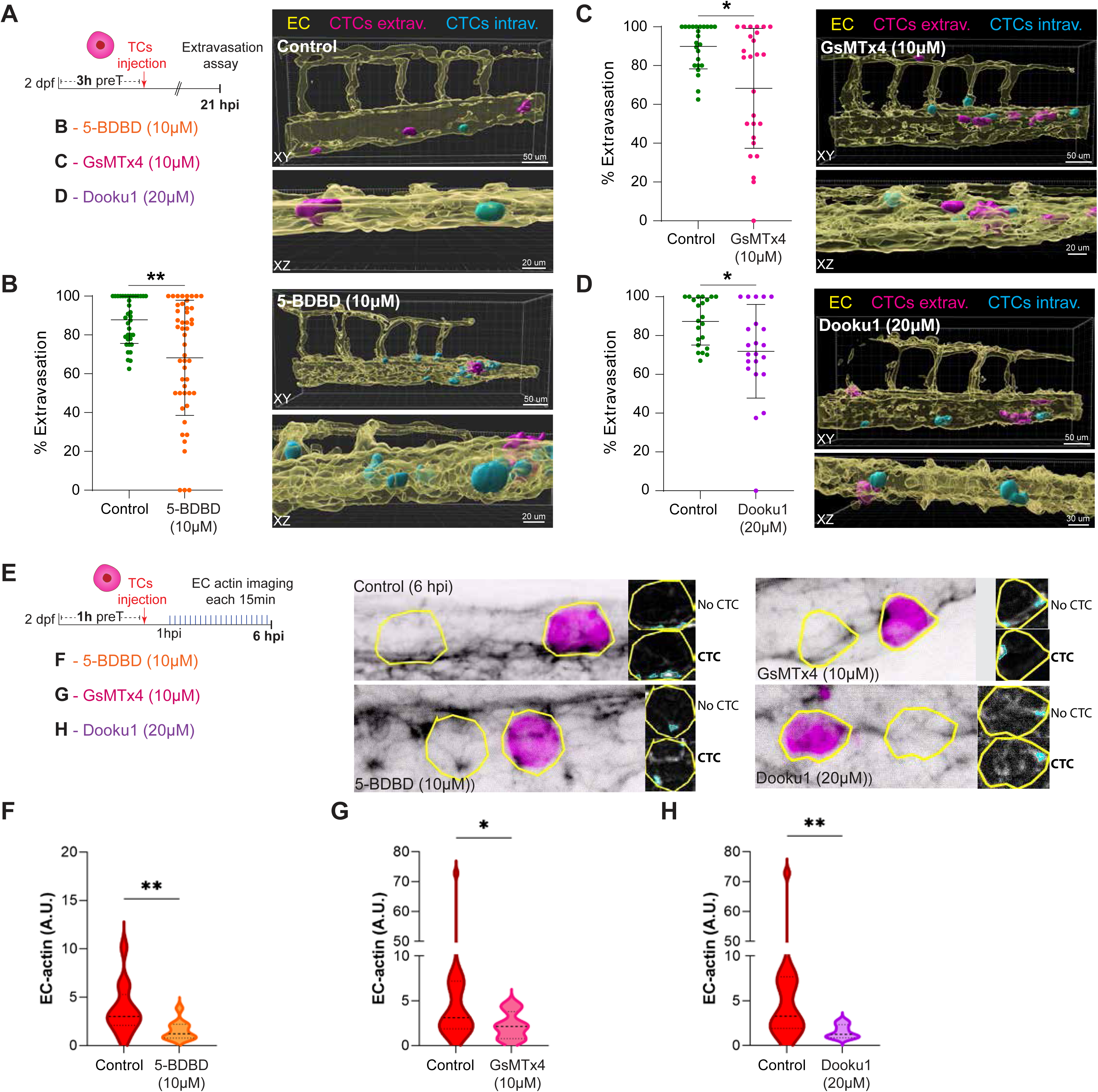
Impairing metastatic extravasation by targeting endothelial mechano-gated calcium channels. **A.** Schematic representation of the experimental design for B, C and D. **B.** Graph shows the percentage of CTC extravasation in control and 5-BDBD-treated embryos at 21 hpi (Mann Whitney test, p-value 0.0018; controls: 37 embryos; 5-BDBD: 48 embryos). Right panels: 3D projections displaying representative examples of control (top) and 5-BDBD- treated embryos (bottom). Transparent EC channel (yellow) facilitates the visualization of intravascular (cyan) and extravascular (magenta) CTCs. **C.** Graph shows the percentage of CTC extravasation in control and GsMTx4-treated embryos at 21 hpi (Mann Whitney test, p-value 0.0018; controls: 24 embryos; GsMTx4: 26 embryos). Right panel: 3D projections displaying representative examples of GsMTx4-treated embryos. Transparent EC channel (yellow) facilitates the visualization of intravascular (cyan) and extravascular (magenta) CTCs. **D.** Graph shows the percentage of CTC extravasation in control and Dooku1-treated embryos at 21 hpi (Mann Whitney test, p-value 0.018; controls: 22 embryos; Dooku1: 22 embryos). Right panel: 3D projections displaying representative examples of Dooku1-treated embryos. Transparent EC channel (yellow) facilitates the visualization of intravascular (cyan) and extravascular (magenta) CTCs. **E.** Schematic representation of the experimental design for F, G and H. **F.** Graph shows EC actin clustering for control (CTC ROIs from non-treated embryos) and 5-BDBD-treated embryos throughout the time lapse in arbitrary units (A.U.) (Mann Whitney test, p value 0.007; control: 6 embryos, 13 CTC-ROIs measured; 5-BDBD: 5 embryos; 11 CTC-ROIs measured for 6 hpi time lapse). **G.** Graph shows EC actin clustering for control (CTC ROIs from non-treated embryos) and GSMTx4-treated embryos throughout the time lapse in arbitrary units (A.U.) (Mann Whitney test, p value 0.0364; control: 11 embryos, 18 CTC- ROIs measured; GSMTx4: 9 embryos; 15 CTC-ROIs measured for 6 hpi time lapse). **H.** Graph shows EC actin clustering for control (CTC ROIs from non-treated embryos) and Dooku1-treated embryos throughout the time lapse in arbitrary units (A.U.) (Mann Whitney test, p value 0.0034; control: 7 embryos, 13 CTC-ROIs measured; GSMTx4: 5 embryos; 7 CTC-ROIs measured for 6 hpi time lapse).

Cancer cells are able to locally secrete cytokines but also act as a direct ATP source to affect their microenvironment. We hypothesized that recently arrested D2A1 cells could secrete factors promoting endothelial remodeling. We performed RNA sequencing of D2A1 cells (Fig. S4) and made several observations. First, D2A1 cells secrete several MMPs including MMP9 that we previously implicated in endothelial remodeling during brain metastasis^25^. These cells also secrete several factors involved in angiogenesis including VEGF-A, ANGPTL2, TGFβ, BMP1 and members of the Notch pathway^60,61^. IGFBP4, the most secreted factor, has also been involved in angiogenesis^62^. Finally, we observed that D2A1 cells have a low expression of ATP secreting channels such as pannexin 1 or connexins, discarding a paracrine effect of arrested CTCs.

As both GsMTx4 and Dooku directly affects Piezo mechano-gated channels and had a strong effect on CTC-induced actin clustering and extravasation, we hypothesized that they might be central players to Ca^2+^ firing induced endothelial remodeling. It has been previously shown that tumor cells overexpress several Ca^2+^ channels including Piezo^58^. To assess whether Ca^2+^ channels from CTCs are involved in extravasation, we depleted Piezo 1, the only Piezo channel expressed in D2A1 cells, and monitored their extravasation efficiency. We did not observe any decrease in extravasation efficiency of Piezo-depleted cells (Fig. S5). This suggests that the role of Piezo mechano-gated channels in CTC extravasation is very likely independent of cancer cells. Ca^2+^ signaling is directly linked to actomyosin contraction through the calmodulin/MLCK pathway^59^. We thus relied on the ML-7 inhibitor which prevents actomyosin contractility by inhibiting MLCK. We observed that ML-7 inhibited CTC-induced endothelial actin clustering (Fig. 5A) and endothelial-driven extravasation (Fig. 5B). Altogether, this suggests that upon adhesion, CTCs modulate mechano-gated calcium channels such as TRPs and/or Piezo, together with P2X4, to induce a Ca^2+^ mechanosignaling, which is transduced through a calmodulin/MLCK pathway to actomyosin remodeling and subsequent endothelial-driven extravasation (Fig. 5C) .

**Figure 5:**
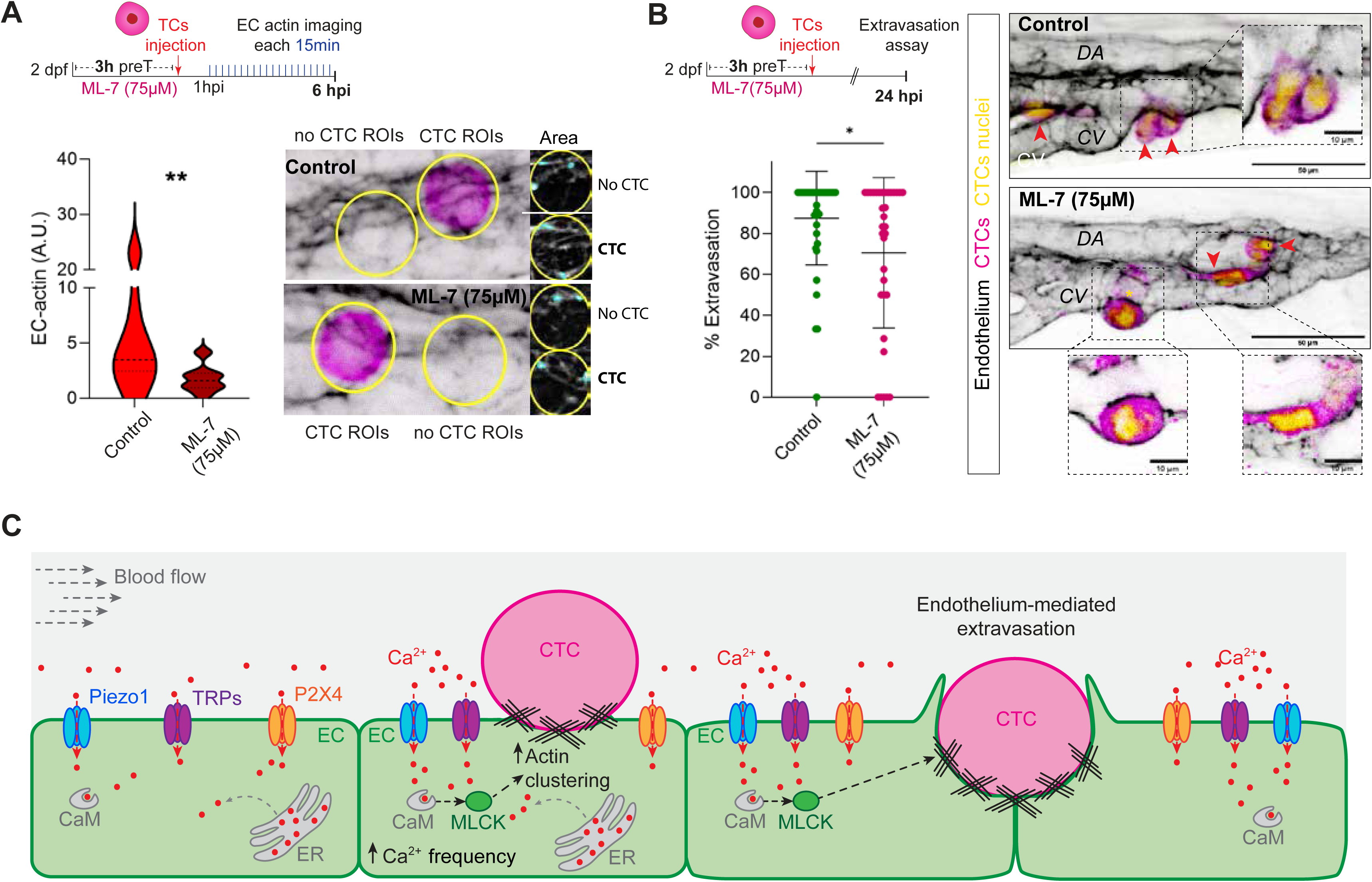
*Impairing Ca^2+^/Calmodulin/MLCK/myosin signaling prevents metastatic extravasation.* **A.** Schematic representation of the experimental design. Graph shows EC actin clustering for control (CTC ROIs from non-treated embryos) and ML-7-treated embryos throughout the time lapse in arbitrary units (A.U.) (Mann Whitney test, p value 0.002; control: 5 embryos, 10 CTC-ROIs measured; ML-7: 8 embryos; 14 CTC-ROIs measured for 6 hpi time lapse). Images show representative examples of the analysis pipeline selections for CTC- and no-CTC- ROIs (yellow line) at 6 hpi for each condition. **B.** Schematic representation of the experimental design. Graph shows the percentage of CTC extravasation in control and ML-7-treated embryos at 21 hpi (Mann Whitney test, p- value 0.0349; control: 40 embryos; ML-7-treated: 34 embryos). Confocal z-stack projections displaying representative examples of control and nifedipine-treated embryos. EC channel is displayed using inverted LUT to facilitate visualization. Zoom boxes show a single confocal plane to improve visualization of intravascular (yellow asterisks) and extravascular (red arrowhead) CTCs. **C.** Schematic representation of the mechanochemical Ca2+ signaling driving extravasation.

## DISCUSSION

In order to successfully form metastasis, tumor cells indoctrinate multiple cell types changing their behavior to facilitate the journey from the primary tumor through the circulation and up to distant colonized organs^63^. Understanding how these indoctrinated behavioral changes occur holds promise to new therapeutic avenues. Here, for the first time, we identify how CTCs indoctrinate ECs into promoting their extravasation. More specifically, we show that CTCs induce an increase in endothelial Ca^2+^ oscillation frequency which triggers the remodeling of the actomyosin cytoskeleton, thus promoting endothelial remodeling—mediated extravasation.

In addition, we provide insight into the cellular dynamics experienced by the indoctrinated endothelium and the molecular players involved. Whereas some CTCs extravasate by diapedesis with little EC contribution, the majority of CTCs we observed, including the highly metastatic clusters, indoctrinate ECs triggering endothelial remodeling to facilitate the process. Recent work shows similarly that Piezo1-mediated Ca^2+^ influx is required for efficient leukocyte diapedesis through MLCK-driven actomyosin reorganization^64^. While this mechanism is very similar to what we observed, the main difference lies in that vascular junctions are maintained throughout endothelial remodeling. Yet, we demonstrate that a similar signaling pathway is involved. This suggests that differences either in the adhesion repertoire or cellular mechanics might account for selecting different extravasation routes. Further work is needed to understand what leads this endothelial plasticity towards diapedesis or remodeling.

We identified Ca^2+^ signaling as the driver of the complex cytoskeletal dynamics undergone by indoctrinated ECs. During angiogenesis, tip cells display increased endothelial Ca^2+^ oscillations that prompt their migratory phenotype^45^. The angiogenic migratory phenotype is characterized by a highly dynamic actin cytoskeleton with migrating tip cells displaying both lamellipodia and filopodia^65,66^. We previously demonstrated that hemodynamic forces induce an angiogenic-like switch in endothelial cells *in vitro* associated with an increased expression of several actin-related genes^24^. Our data revealed how the presence of CTCs increased the frequency of endothelial Ca^2+^ oscillations, promoting a cytoskeletal reorganization that allows ECs to alter the vascular integrity and actively participate in the extravasation process. We observed that early on after CTC arrest (2 hpi), Ca^2+^ oscillations trended to be increased closer to arrested CTCs, which suggests that TCs can indoctrinate EC signaling locally, either by direct interaction or by short-range paracrine signaling. Interestingly, later in the process (6 hpi), all ECs displayed increased oscillatory frequency, suggesting that ECs synchronized their Ca^2+^ signaling over time. These Ca^2+^ oscillation have been suggested to reflect VEGFR signaling^45^. Since we observed that flow did not influence Ca^2+^ oscillations in our model, it is tempting to speculate that VEGFR2-flow-induced signaling acts downstream of Ca^2+^ flux. However, as we are relying on a highly diffusible genetically encoded probe expressed in the whole cell, we cannot exclude that that it might not be sensitive enough to track Ca^2+^ firing at subcellular levels that could be produced by subtle flow changes. We have shown that nifedipine also affects WM983B melanoma extravasation, though less efficiently. This supports the idea that extravasation depending on endothelial Ca^2+^ firing is not specific to D2A1 cells, in accordance with the literature which showing that the human breast cancer line MDA-MB-231 and the mouse breast cancer line E0771^23,25^ also extravasate through endothelial remodeling. This suggests that cell specific mechanisms are also involved. Interestingly, neither inert nor FN-coated beads were able to provoke calcium changes engaging endothelial remodeling with the same magnitude as CTCs. This might suggest that CTCs are very likely pre-activating ECs through direct adhesion engagement and/or secreted factors. The biochemical cues and the adhesion molecules that CTCs leverage upon ECs, whether they are cancer-type specific, might account for their organ tropism and require further research.

The pharmacological approach we relied on is very likely affecting CTCs as well. D2A1 cells express ROCK1 and 2 but also Ca^2+^ channels including Piezo1/2, TRPs (C2/4, TRPM2-8 and V2/4) P2X3-7 and P2Y12/14 and L-type channels Cav1.2 (unpublished data). Besides, calcium signaling was shown to alter CTC cytoskeletal remodeling, thus affecting their metastatic capabilities^67–69^. We cannot rule out that the extravasation reduction observed upon calcium inhibition by nifedipine might result from the synergic action on CTCs and ECs. However, in a patient context, any systemic therapeutic approach would target both CTCs and the endothelium. We believe our present work suggests that the decrease in extravasation observed upon Ca^2+^ inhibition could be exploited by treating patients subjected to primary tumor resection where CTC number is increased to reduce the metastatic risk.

## AUTHOR CONTRIBUTIONS

Conceptualization: M.P. with inputs from N.O. and J.G.G.; Endothelial actin imaging: M.P. and A.D.; Calcium imaging: M.P; Extravasation assays: M.P. and A.D.; cell engineering: O.L. and A.L., RNAseq: T.S., A.M. and R.C., beads production and characterization: R.G. and S.G.; Data analysis: M.P. and A.D.; Project managing: M.P., N.O. and J.G.G.; Writing manuscript: N.O, M.P. and J.G.G. Funding acquisition: N.O. and J.G.G.

## ACKNOWLEDGMENTS

We are grateful to Julien Vermot for the *Tg(fli:GalFF; UAS:GCAMP7a)*, *Tg(fli:lifeAct-eGFP)* and *Tg(fli:lifeAct-eGFP;flk:nls-mCherry)* and Holger Gerhardt the *Tg(fli:EGFP-CAAX)* transgenic lines kindly provided for this paper, Sara Jimenez for her very valuable contribution to optimizing the image analysis used, and Sebastien Harlepp for fruitful discussions on calcium analysis, PICSTRA (CRBS, Pascal Kessler) and IGBMC microscopy facilities and thank Parth Patel, part of TDSU Lab-on-a-chip systems at MPL, for the production of the master template and microfluidic chips used for the bead production. We thank members of the Tumor Biomechanics lab for helpful discussion and particularly Vincent Hyenne, Nandini Asokan and Zeynep Yesilata for preliminary data.

## STAR METHODS

### KEY RESSOURCES TABLE

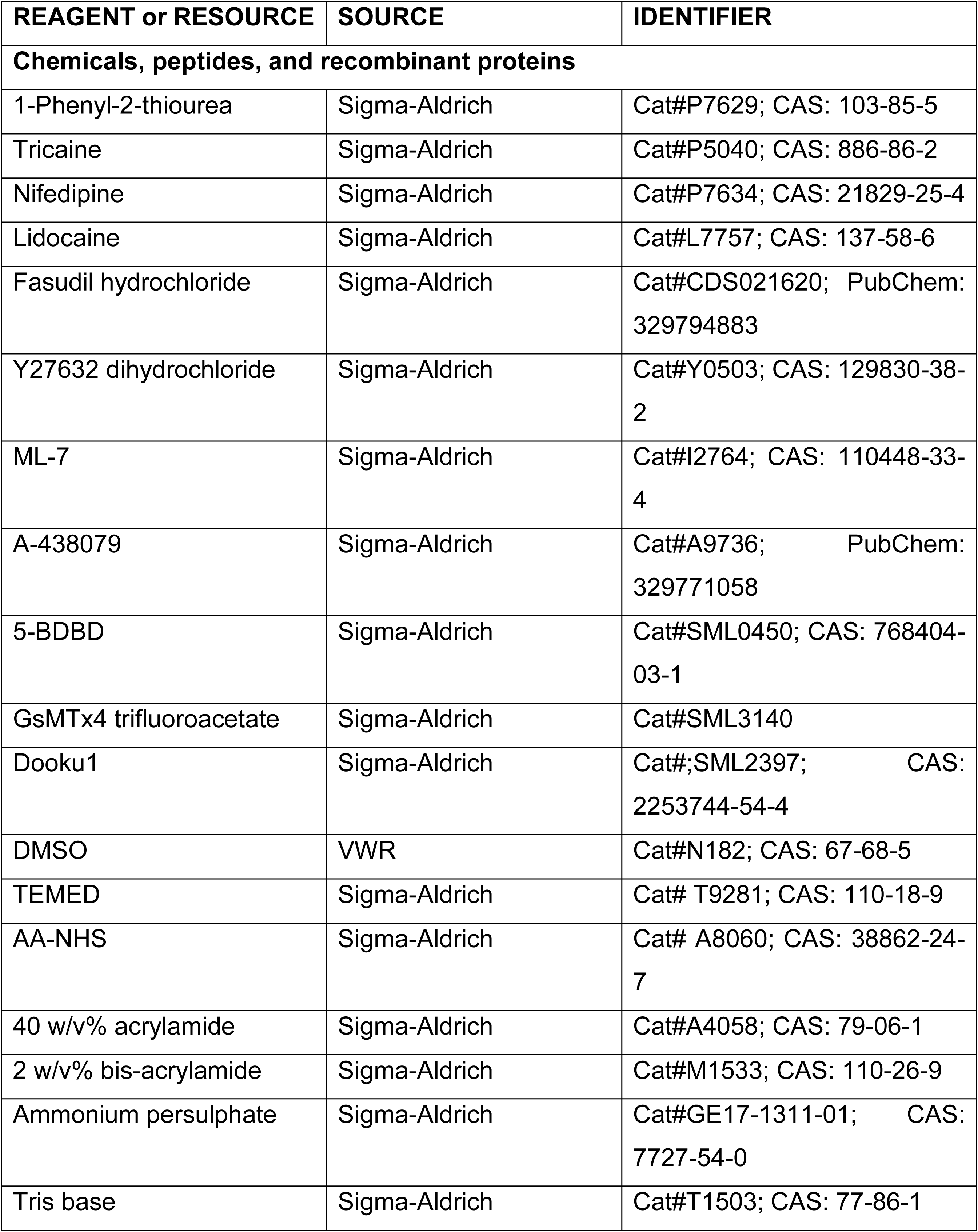
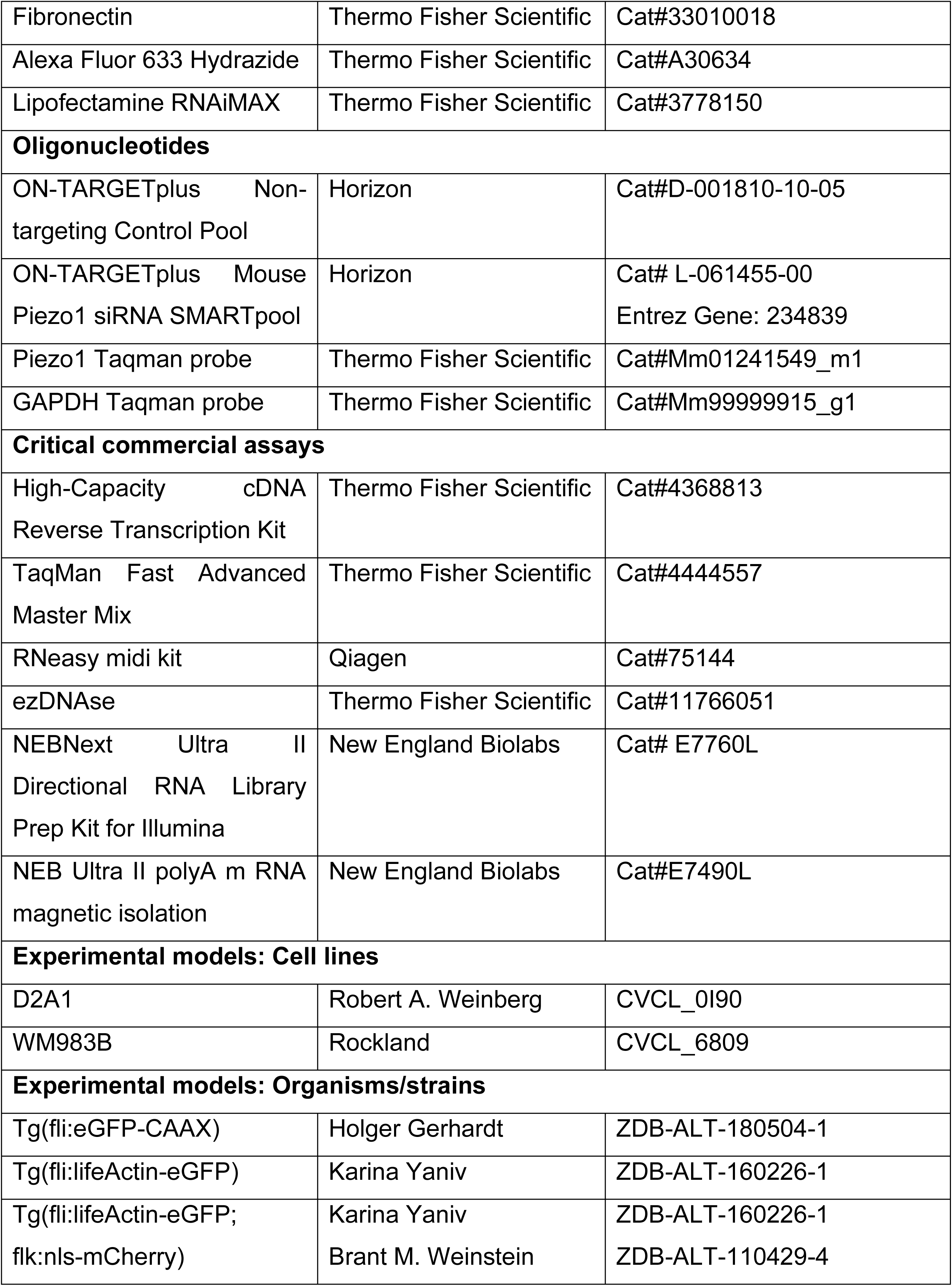
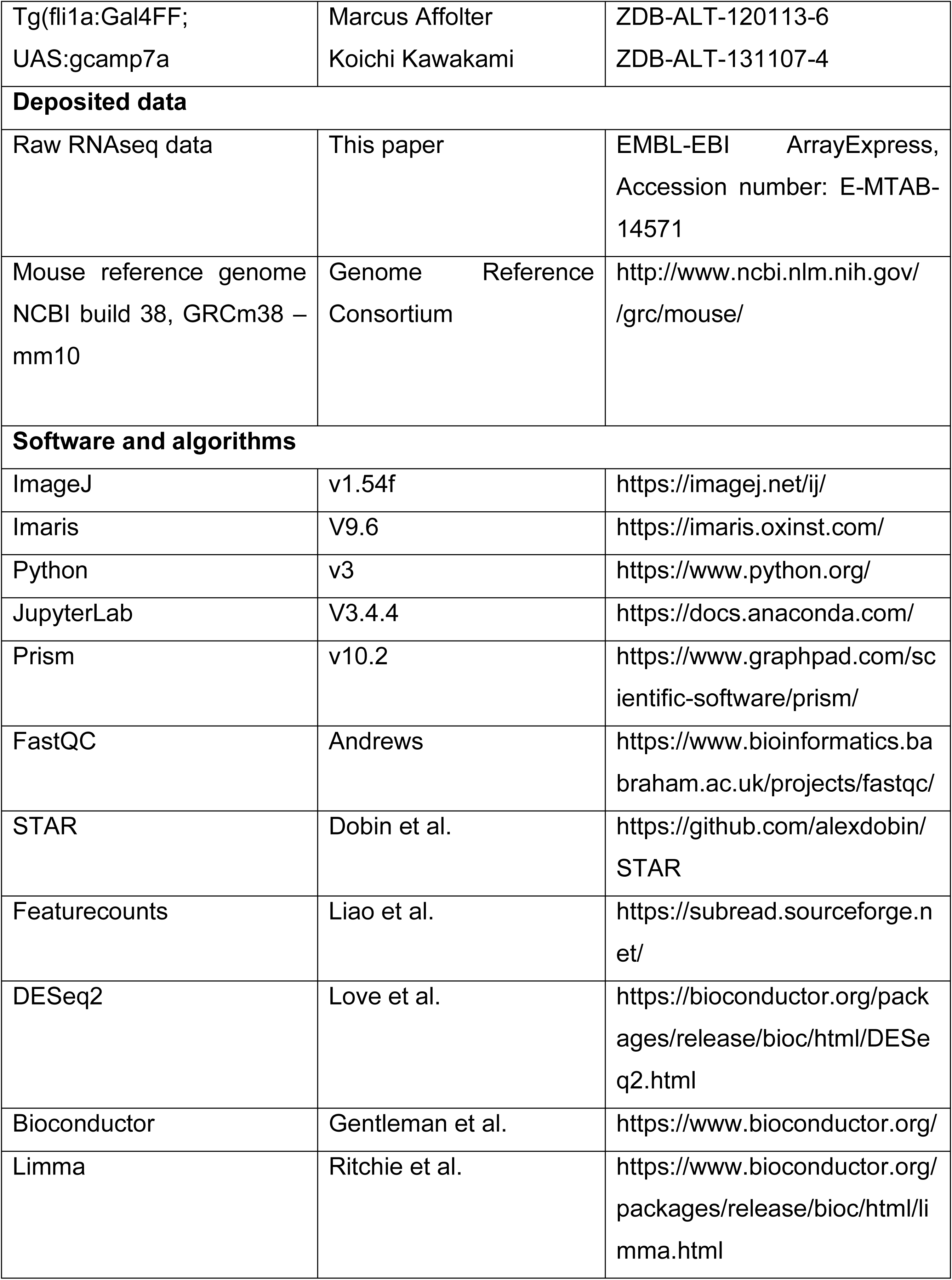

### EXPERIMENTAL MODEL

#### Cell lines

D2A1 native cells were kindly provided by Robert A. Weinberg (MIT). Cells were grown in DMEM with 4.5 g/l glucose (Dutscher) supplemented with 5% FBS (Dutscher), 5% NBCS (Thermo Fischer Scientific), 1% NEAA (Thermo Fischer Scientific) and 1% penicillin-streptomycin (Dutscher). Cells were transfected with ONTARGETplus siRNAs (Horizon) using Lipofectamine RNAiMAX (Thermo Fischer Scientific) following the manufacturer’s instructions. Experiments were performed 72 h post-transfection.

WM983B is a patient-derived human metastatic melanoma cell line mutant for BRAF V600 and was purchased from Rockland. Cells were cultured in MCDB153 (Dutscher) and Leibovitz’s L-15 medium (Thermo Fisher Scientific) in a 4 to 1 ratio, supplemented with 2% foetal bovine serum, 1.68 mM CaCl2 and 1% penicillin/streptomycin (Dutscher). Cells stably expressing LifeAct-TdTomato and LifeAct-miRFP were obtained in house using lentiviral transduction.

#### Zebrafish handling and drug treatment

*Tg(fli:lifeActin-eGFP), Tg(fli:eGFP-CAAX), Tg(fli:lifeActin-eGFP;flk:nls-mCherry)* and *Tg(fli1a:Gal4FF; UAS:gcamp7a)* zebrafish (*Danio rerio*) embryos were maintained in Danieau 0.3X medium (17,4 mM NaCl, 0,2 mM KCl, 0,1 mM MgSO4, 0,2 mM Ca(NO3)2) buffered with HEPES 0,15 mM (pH = 7.6), supplemented with 200 µM of 1-Phenyl-2- thiourea (PTU, Sigma-Aldrich) to inhibit melanogenesis, as previously described^70^. Nifedipine (5μM in DMSO, Sigma-Aldrich) or lidocaine (0.015%, Sigma-Aldrich) were added in water (Danieau 0.3X + PTU) of the embryos 1 hour (h) before injection of tumor cells and maintained throughout the experiments. Fasudil hydrochloride (30μM in DMSO, Sigma-Aldrich), Y27632 dihydrochloride (45μM in water, Sigma-Aldrich), ML-7 (75µM in DMSO, Sigma-Aldrich), 5-BDBD (10μM in DMSO, Sigma-Aldrich), A-438079 (10μM in DMSO, Sigma-Aldrich), GsMTx4 trifluoroacetate (10μM in water, Sigma-Aldrich) or Dooku1 (20μM in DMSO, Sigma-Aldrich) were added in water (Danieau 0.3X + PTU) of the embryos 3 hours (h) before injection of tumor cells and maintained throughout the experiments. All zebrafish procedures were performed in accordance with French and European Union animal welfare guidelines and supervised by local ethics committee (ZF facility A6748233; APAFIS 2018092515234191).

### METHOD DETAILS

#### Production of polyacrylamide microgel beads

Polyacrylamide microgel beads were produced as previously described by Girardo et al.^71^. In brief, microemulsions of a polyacrylamide pre-gel mixture in fluorinated oil (3MTM NovotecTM 75000, Iolitec Ionic Liquids Technologies GmbH, Germany) were generated using a droplet microfluidic platform. The fluorinated oil contained 2.4 w/v% ammonium Krytox® surfactant, 0.4 v/v% N, N, N′, N′-tetramethylethylenediamine (TEMED, Sigma-Aldrich) as the catalyst, and 0.1% v/v acrylic acid N-hydroxysuccinimide-acrylamide ester (AA-NHS, Sigma-Aldrich) to facilitate the binding of fluorophores and proteins. The pre-gel mixture comprised 40 w/v% acrylamide (Sigma-Aldrich) as the monomer, 2 w/v% bis-acrylamide (Sigma-Aldrich) as the crosslinker, and 0.05 w/v% ammonium persulphate (Sigma-Aldrich) as the free radical initiator, diluted in 10mM Tris-buffer (pH = 7.48). To make the beads fluorescent, Alexa Fluor 633 Hydrazide (3 mg/ml), (Invitrogen) solution was added to the pre-gel mixture (1.8 v/v%). For the FN-coated beads, the produced beads underwent washing in a 50mM HEPES solution (pH 8.22) and approximately 10 million beads were incubated with 200 µl FN solution (0.5 mg/ml), (Thermo Fisher Scientific) for 48 h at 4 °C on a vertical rotator at 20 rpm. The final beads were washed 3 times in 1X PBS and stored at 4 °C. Fine-tuning of the final size and elasticity of the beads was achieved by adjusting the flow conditions on the microfluidic chip and the total monomer concentration in the pre-gel mixture. Bead size distribution was determined by analyzing bright-field images using a macro implemented in freeware Fiji. Additionally, the Young’s modulus of the beads was measured by real-time deformability cytometry^72^. Three distinct batches of beads were produced, with the following values for diameters (*dmean* ± SD) and Young’s moduli (*Emean* ± SD):

- Inert polyacrylamide microgel beads: *d* = (15.2 ± 0.5) µm, *E* = (1.6 ± 0.3) kPa
- FN-coated microgel beads: *d* = (16.5 ± 0.9) µm and *E* = (0.5 ± 0.1) kPa; d = (16.5 ± 0.7) µm and E = (1.9 ± 0.3) kPa

#### Intravascular injection

48-hour post-fertilization (hpf) embryos were mounted in 0.8% low melting point agarose pad containing 650 µM of tricaine (ethyl-3-aminobenzoate-methanesulfonate) to immobilize the embryos. D2A1 cells were injected with a Nanoject microinjector 2 (Drummond) and glass capillaries (25 to 30 µm inner diameter) filled with mineral oil (Sigma). 13.8nL of cell suspension at 100.10^6^ cells per ml were injected in the duct of Cuvier of the embryos under the M205 FA stereomicroscope (Leica). Polyacrylamide microgel beads at a respective stock concentration of 1.6kPa-Alexa Fluor 633 at 83 x 10^6^ beads/ml); 1.9kPa-fibronectin-coated Alexa Fluor 633 at 73 x 10^6^ beads/ml or 0.5kPa- fibronectin-coated Alexa Fluor 633 at 105 x 10^6^ beads/ml) were diluted to 2μl beads in 7μl Danieau, then injected using the same equipment.

#### EC actin clustering analysis

Confocal imaging was performed with an inverted Olympus IX83 with a CSU-W1 spinning disk head (Yokogawa), an ORCA Fusion Digital camera, and a UPL SAPO 30x 1.05 NA silicone immersion objective. Depending on the experiment, the caudal plexus region of embryos treated or untreated after D2A1 or polyacrylamide microgel beads injections was imaged with a z-step of 3µm every 15 min for 4 or 6 hpi starting from 1 hpi. A minimum of 3 embryos per condition from at least 3 independent experiments. A minimum of 10 CTCs per condition were analyzed. Image analysis was performed using ImageJ (https://imagej.nih.gov/ij/index.html). Z-stack sum projections were made for each time point. We manually draw regions of interest (ROIs) around individual CTCs through time (single CTCs or clusters of 2 cells) and identical ROIs in equivalent regions lacking CTCs. Top Hat filter, radius=10, was applied to the ROIs, and actin clusters were selected using MaxEntropy dark thresholding and running Analyze Particles, size = 1 - infinity. We calculated the percentage of EC actin clustering, i.e., area of actin clustered divided by the total area of the ROI measured, for each pair of ROIs (CTC and no-CTC) throughout the time-lapse. Afterward, we calculated the area under the curve (AUC) for all the time points in the time-lapse for each set of ROIs (CTC and no-CTC), which provided us with a cumulative measurement of actin dynamics through time. Lastly, we normalized each CTC-AUC measurement to its no-CTC-AUC control (Fig.1B). We validated the analysis pipeline specificity by drawing two no-CTC ROIs for each time point in typical arrest regions of the vasculature and assigning them randomly to group A or B. Then, we performed the previously described analysis normalizing group A to B and conversely group B to A (Fig.S1B and D).

#### EC Ca^+2^ analysis

Confocal imaging was performed with an inverted Nikon TI Eclipse with a CSU-X1 spinning disk head, dual camera Photometrics Prime 95B, and a 20x 0.75 NA oil immersion objective. A single plane of the caudal plexus region of each embryo was imaged at 200 ms for 5 minutes (1 500 frames in total). Image analysis was performed using ImageJ. We applied the Bleach correction (exponential fit) to the z-stack, calculated the average intensity projection, and subtracted the average intensity projection from the bleach-corrected z-stack, to highlight changes in GFP intensity. Manually, we selected EC cells that displayed calcium oscillations and extracted the oscillations amplitude and frequency using “Plot z-axis profile”.

#### Extravasation analysis

Confocal imaging was performed with an inverted Olympus IX83 with a CSU-W1 spinning disk head (Yokogawa), an ORCA Fusion Digital camera, and an UPL SAPO 30x 1.05 NA silicone immersion objective. The caudal plexus region of embryos treated or untreated after D2A1 or polyacrylamide microgel beads injections with a z-step of 3µm. Manual quantification using ImageJ of the extravasated CTCs or polyacrylamide microgel beads over the total to obtain the % of extravasation at 21 hpi.

#### RT-qPCR

Total RNAs were isolated from transfected D2A1 cells using a RNeasy midi kit (Qiagen). After total RNA ezDNAse treatment (Thermo Fisher Scientific), cDNAs were obtained using High-Capacity cDNA Reverse Transcription Kit (Thermo Fisher Scientific). RT-qPCR reactions were made using TaqMan Fast Advanced Master Mix (Thermo Fisher Scientific) in a CFX Connect Real-Time PCR Detection System (Biorad). Amplification results were normalized using GAPDH levels and relative expression evaluated using the ΔΔcT method^73^.

#### RNAseq

RNA integrity was assessed by Bioanalyzer (total RNA Pico Kit, 2100 Instrument, Agilent Technologies, Paolo Alto, CA, USA). All samples had RNA integrity numbers above 8 and DV200>80%. Sequencing libraries were prepared using “NEBNext Ultra II Directional RNA Library Prep Kit for Illumina“ and enriched in mRNA using “NEB Ultra II polyA mRNA magnetic isolation” kit (New England Biolabs, Ipswich, MA, USA). Libraries were pooled and sequenced (single-end, 100bp) on a NextSeq2000 according to the manufacturer’s instructions (Illumina Inc., San Diego, CA, USA). For each sample, quality control was carried out and assessed with the NGS Core Tools FastQC (Andrews et al., 2010, http://www.bioinformatics.babraham.ac.uk/projects/fastqc). Sequence reads (minimum 48,5 million) were mapped to Mus musculus mm10 using STAR^74^ to obtain a BAM (Binary Alignment Map) file. An abundance matrix was generated based on read counts identified by Featurecounts^75^. At last, differential expression analyses were performed using the DESeq2^76^ package of the Bioconductor framework for RNASeq data^77^. Up and down- regulated genes were selected based on the adjusted p-value (< 0.05) and the fold-change (> 1.5). Raw RNAseq data have been deposited in the EMBL-EBI ArrayExpress archive (accession number E-MTAB-14571).

Multidimensional scaling (MDS) was performed on gene expression counts normalized using the DESeq2 R package^76^. MDS coordinates were calculated using the plotMDS function from the limma R package^78^. This function calculates pairwise distances between samples based on their expression profiles. The top two dimensions were retained for visualization, representing the major sources of variation in the dataset, typically reflecting biological or technical differences between samples.

#### Statistical analysis

Statistical analysis was performed using the GraphPad Prism version 10 software. The Shapiro-Wilk normality test was used to confirm the normality of the data. A Student unpaired two-tailed t-test was used for data following a Gaussian distribution. For data not following a Gaussian distribution, the Mann-Whitney test was used. When more than 3 groups were compared, a Kruskal-Wallis test followed by Dunn’s Multiple Comparison test was used. The Wilcoxon matched-pairs rank test was used for EC actin analysis of CTC-ROIs and its no-CTC-ROIs control. Illustrations of these statistical analyses are displayed as the mean +/- standard deviation (SD). P-values smaller than 0.05 were considered as significant. *, p<0.05, **, p<0.01, ***, p<0.001, ****, p<0.0001.

## SUPPLEMENTAL FIGURE LEGENDS

**Figure S1:**
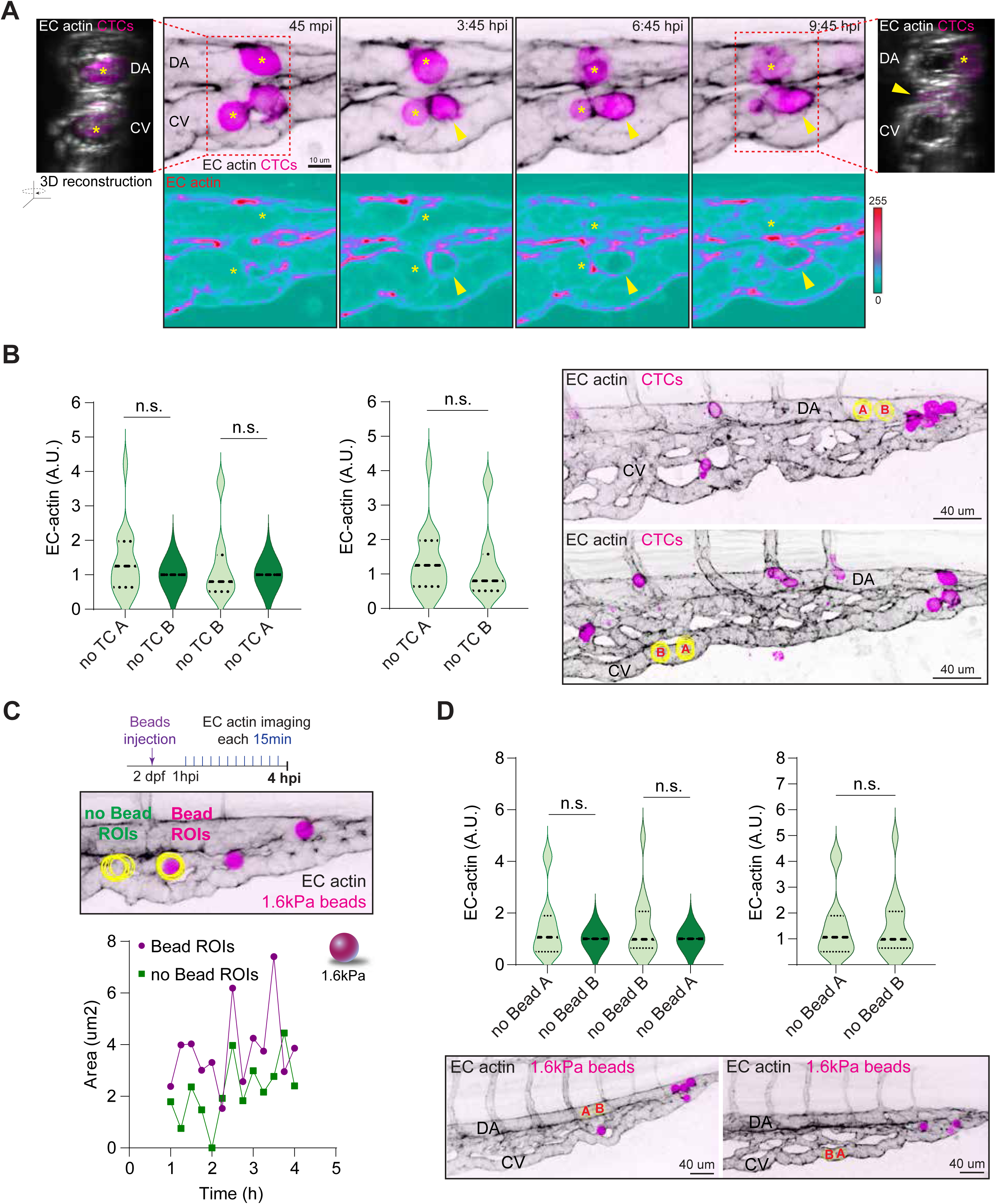
Endothelial actomyosin dynamics mediate CTC extravasation. **A.** Time-lapse z-stack projections show CTCs extravasating from the DA and CV. 3D orthogonal views at time points 45 mpi (top left) and 9:45 hpi (top right) facilitate intravascular and extravascular CTC visualization. Bellow, individual actin channel is displayed using ice LUT to facilitate the visualization of signal intensity, where green is the minimum and red is the maximum. For all the images, yellow asterisks label intravascular CTCs, yellow arrowhead mark the extravasated CTC. **B.** Left-side graph shows actin analysis pipeline validation for CTCs where EC-actin was quantified for two sets of equivalent no-CTC ROIs (A and B), normalized to each other. No differences appeared using each one as control (Wilcoxon matched-pairs signed rank test, no TC A normalized to B: p-value 0.33; no TC B normalized to A: p-value 0.97, 4 embryos, two sets of 17 no-CTC- ROIs measured for 11 hpi time-lapse). Right-side graph shows the comparison between the two set of normalized EC-actin quantifications without differences. These results highlighted the specificity of the EC-actin dynamics triggered by CTCs (Mann Whitney test, p-value 0.4, 4 embryos, two sets of 17 no-CTC- ROIs measured for 11 hpi time-lapse). Right-side images show examples of no-CTC equivalent ROIs randomized in groups A or B (yellow circles) projected over the first time point of the time-lapse. **C.** Schematic representation of the experimental design. Example of EC- actin quantification for 1.6kPa beads: Yellow ROIs (1.6kPa bead and no-bead) are projected over the first time point of the time-lapse. Graph bellow shows EC-actin area through time. **D.** Left-side graph shows actin analysis pipeline validation for 1.6kPa beads where EC-actin was quantified for two sets of equivalent no-bead ROIs (A and B), normalized to each other. No differences appeared using each one as control (Wilcoxon matched-pairs signed rank test, no-beads A normalized to B: p-value 0.26; no-beads B normalized to A: p-value 0.42, 5 embryos, two sets of 22 no-CTC- ROIs measured for 4 hpi time-lapse). Right-side graph shows the comparison between the two set of normalized EC-actin quantifications without differences (Mann Whitney test, p value 0.94, 5 embryos, two sets of 22 no-CTC- ROIs measured for 4 hpi time lapse). Images show examples of no-bead equivalent ROIs randomized in groups A or B (yellow circles) projected over the first time point of the time-lapse.

**Figure S2:**
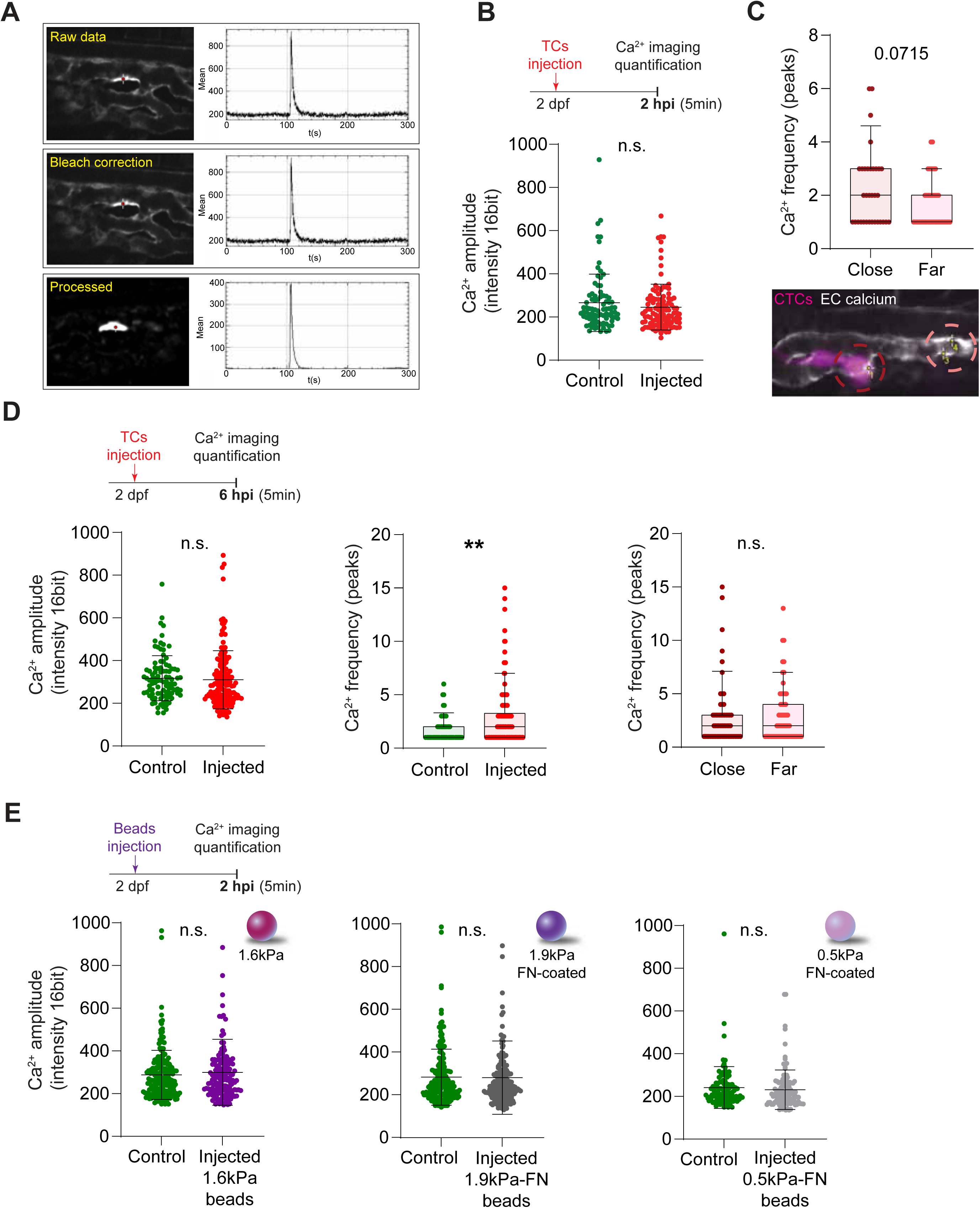
CTC arrest alters EC calcium signaling. **A.** Examples of the EC calcium analysis pipeline. Upper image shows a raw confocal projection displaying an EC calcium firing (red dot). Right-side graph shows the calcium firing quantification. Middle image shows the same EC after bleach correction. Right-side graph shows the calcium firing quantification. Lower image shows the same EC processed (after subtracting the average intensity projection from the bleach corrected z-stack, to highlight changes in GFP intensity). Right-side graph shows the calcium firing quantification. **B.** Schematic representation of the experimental design. Graph shows EC calcium firing amplitude (signal intensity) in non- injected (control) and injected embryos at 2 hpi (Mann Whitney test, p value 0.29; controls: 5 embryos, 84 cells quantified; injected: 5 embryos, 106 cells quantified). **C.** Graph shows EC calcium firing frequency (number of peaks) in injected embryos at 2 hpi. ECs were sorted into two groups: close (in direct contact with CTCs) or far (not in direct contact with CTCs, Mann Whitney test, p-value 0.071; close: 5 embryos, 33 cells quantified; far: 5 embryos, 43 cells quantified). Confocal plane illustrating the close and far categories. Numbered yellow dots exemplified some of the measured ECs. Graph is a boxplot (upper/lower quartile, median, bars show the 10%-90% range), each dot represents an individual EC. **D.** Schematic representation of the experimental design. Left-side graph shows EC calcium firing amplitude (maximum intensity) in non- injected (control) and injected embryos at 6 hpi (Mann Whitney test, p value 0.12; controls: 5 embryos, 95 cells quantified; injected: 6 embryos, 156 cells quantified). Middle graph shows EC calcium firing frequency (number of peaks) in non- injected (control) and injected embryos at 6 hpi (Mann Whitney test, p value 0.0019; controls: 5 embryos, 66 cells quantified; injected: 6 embryos, 126 cells quantified). Right-side graph shows EC calcium firing frequency (number of peaks) in injected embryos at 6 hpi. ECs were sorted into two groups: close (in direct contact with CTCs) or far (not in direct contact with CTCs, Mann Whitney test, p-value 0.9; close: 5 embryos, 58 cells quantified; far: 6 embryos, 68 cells quantified). Frequency graphs are boxplots (upper/lower quartile, median, and bars show the 10%-90% range); each dot represents an individual EC. **E.** Schematic representation of the experimental design. Graphs show EC calcium firing amplitude (signal intensity) in non-injected (control) and injected embryos at 2 hpi. Left-side graph shows 1.6kPa beads injections (Mann Whitney test, p value 0.69; controls: 7 embryos, 185 cells quantified; injected: 8 embryos, 154 cells quantified). Middle graph shows 1.9kPa-fibronectin-coated beads injections (Mann Whitney test, p value 0.327; controls: 8 embryos, 178 cells quantified; injected: 7 embryos, 182 cells quantified). Right-side graph shows 0.5kPa-fibronectin-coated beads injections (Mann Whitney test, p value 0.11; controls: 4 embryos, 100 cells quantified; injected: 5 embryos, 113 cells quantified).

**Figure S3:**
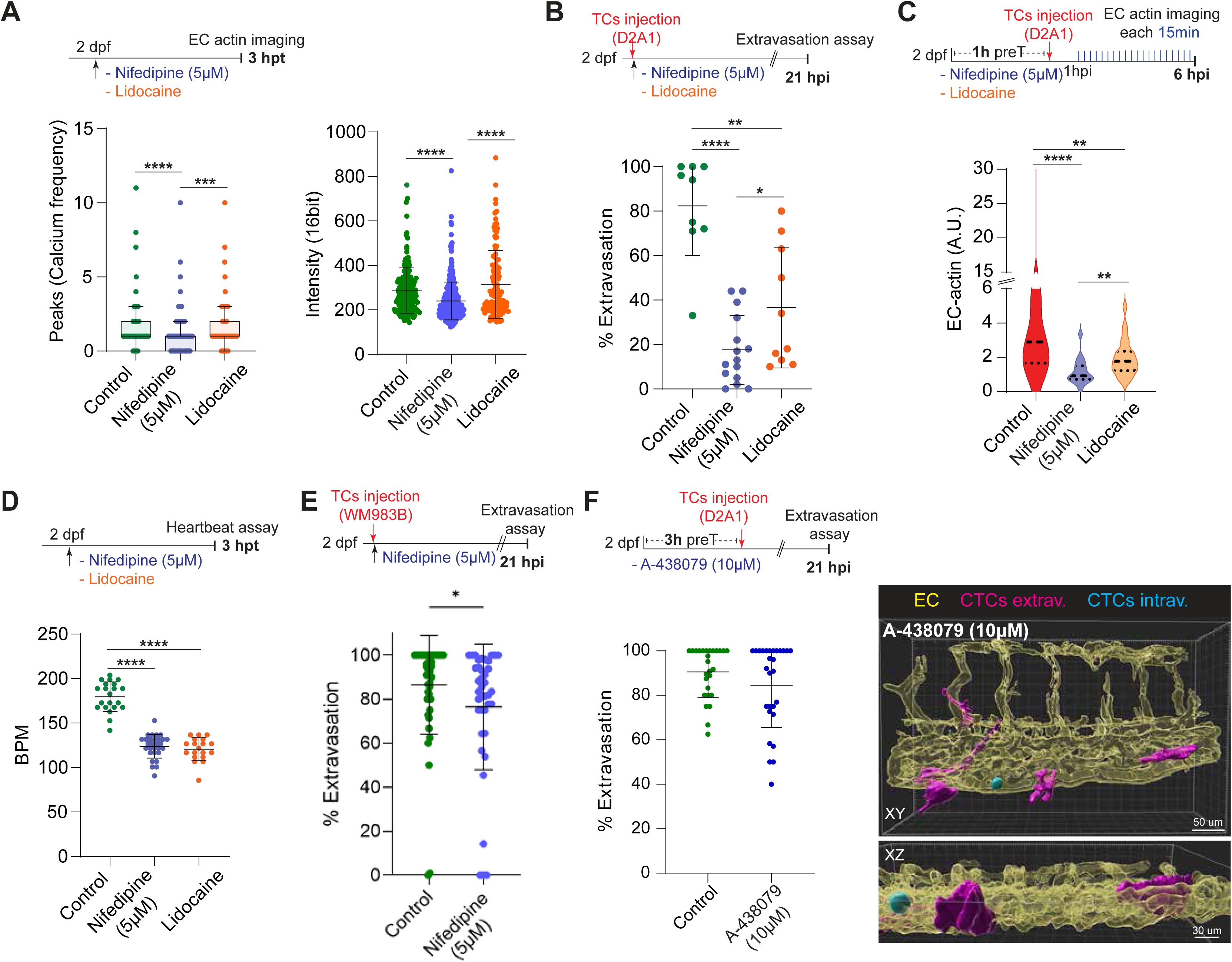
EC mechano-gated channels mediating CTC extravasation. **A.** Schematic representation of the experimental design. Left-side graph shows EC calcium firing frequency (number of peaks) in control (non-treated), nifedipine-, and lidocaine-treated embryos at 3 hours post-treatment (hpt, Kruskal-Wallis test multiple comparison control – nifedipine: p-value <0.00001; control – lidocaine: p-value 0.0006; controls: 7 embryos, 171 cells quantified; nifedipine: 11 embryos, 354 cells quantified; lidocaine: 6 embryos, 120 cells quantified). This graph is a boxplot (upper/lower quartile, median, and bars show the 10%-90% range); each dot represents an individual EC. Right-side graph shows EC calcium firing amplitude (maximum intensity) in control (non-treated), nifedipine-, and lidocaine-treated embryos at 3 hpt (Kruskal-Wallis test multiple comparison control – nifedipine: p-value <0.0001; control – lidocaine: p-value <0.0001; controls: 7 embryos, 171 cells quantified; nifedipine: 11 embryos, 355 cells quantified; lidocaine: 6 embryos, 120 cells quantified). **B.** Schematic representation of the experimental design. Graph shows percentage of CTC extravasation in control, nifedipine- and lidocaine-treated embryos at 21 hpi (Mann-Whitney: control vs nifedipine-treated: p value 0.0001; control vs lidocaine: p value 0.0021; nifedipine vs lidocaine: p value: 0.0341; control: 9 embryos; nifedipine-treated: 15 embryos; lidocaine-treated: 10 embryos). **C.** Schematic representation of the experimental design. Graph shows EC actin clustering for control (CTC-ROIs from non-treated embryos), nifedipine-, and lidocaine-treated embryos throughout the time lapse in arbitrary units (A.U., Control vs nifedipine-treated: Mann Whitney test, p-value 0.0001; control: 3 embryos, 19 CTC- ROIs measured; nifedipine: 2 embryos, 18 CTC- ROIs measured for 6 hpi time-lapse; Control vs lidocaine-treated: Mann Whitney test, p-value 0.009; control: 3 embryos, 19 CTC- ROIs measured; lidocaine: 2 embryos, 25 CTC- ROIs measured for 6 hpi time-lapse; Nifedipine vs lidocaine-treated: Mann Whitney test, p-value 0.003; nifedipine: 2 embryos, 18 CTC- ROIs measured; lidocaine: 2 embryos, 25 CTC- ROIs measured for 6 hpi time-lapse).**D.** Schematic representation of the experimental design. Graph shows heartbeat in beats per minute (BPM) in control, nifedipine-, and lidocaine-treated embryos (ANOVA multiple comparisons: control vs nifedipine p value <0.0001; control vs lidocaine p value <0.0001; control: 21 embryos; nifedipine: 29 embryos; lidocaine: 19). **E.** Schematic representation of the experimental design. Graph shows percentage of CTC extravasation in control and nifedipine-treated embryos at 21 hpi (Mann Whitney test, p value 0.0168; controls: 46 embryos nifedpipine: 39 embryos). **F.** Schematic representation of the experimental design. Graph shows percentage of CTC extravasation in control and A-438079-treated embryos at 21 hpi (Mann Whitney test, p value 0.42; controls: 27 embryos; A-438079: 28 embryos). 3D projections displaying a representative example of an A-438079-treated embryo. Transparent EC channel (yellow) facilitates the visualization of intravascular (cyan) and extravascular (magenta) CTCs.

**Figure S4:**
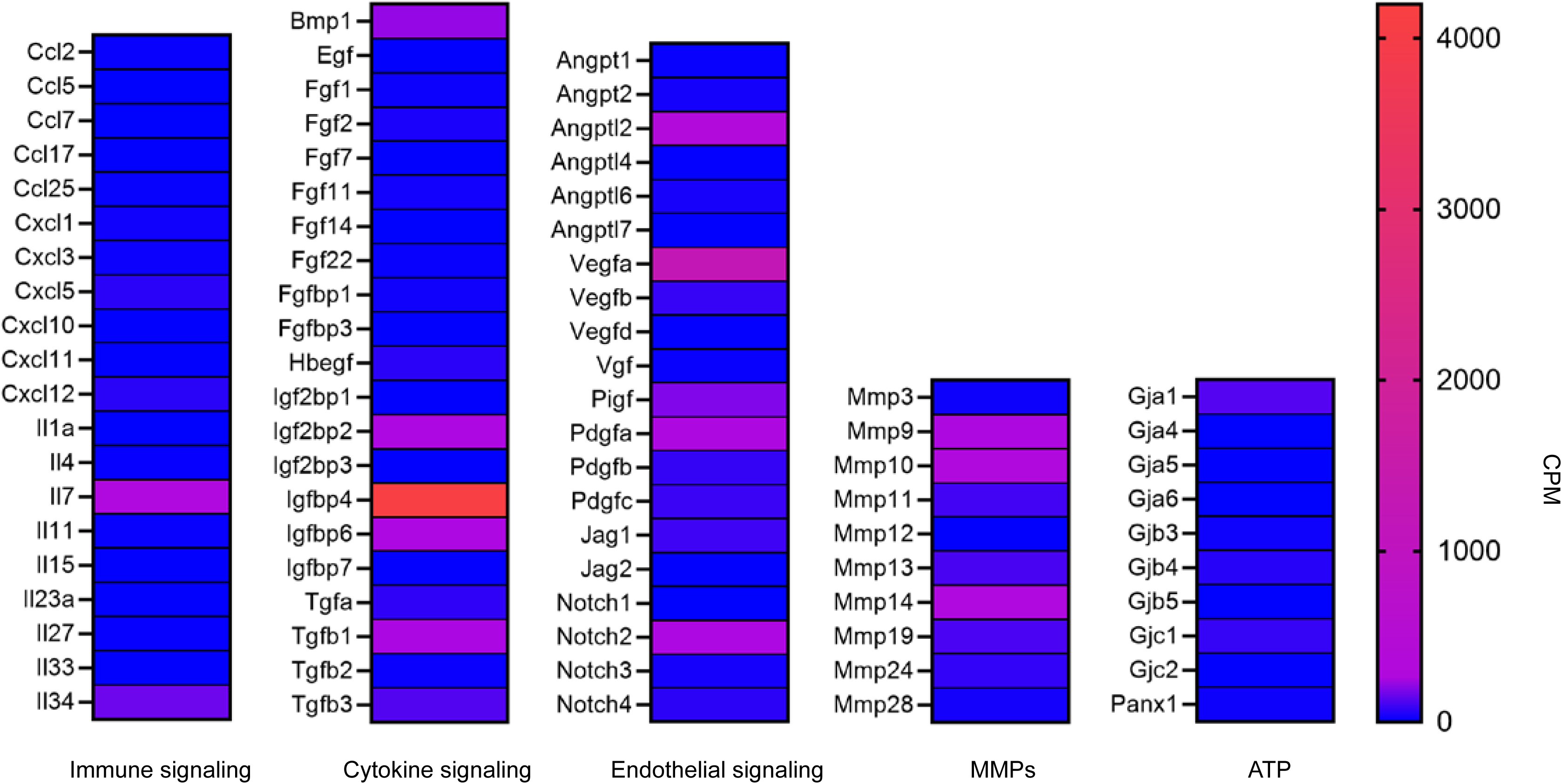
D2A1 cells secrete factors involved in endothelial signaling. D2A1 cells secretome was assessed by RNA sequencing followed with GO analysis. Secreted factors are divided by GO terms; values are displayed in counts per million (CPM).

**Figure S5:**
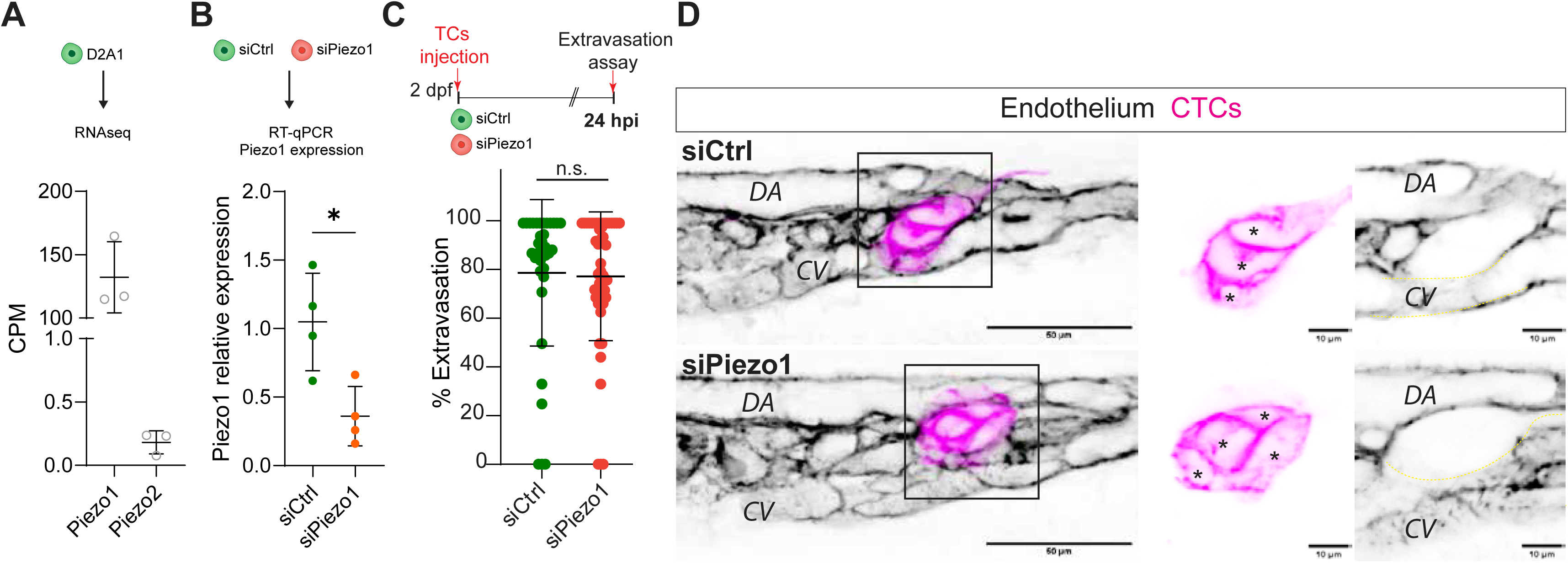
CTC Piezo mechano-gated are not involved in extravasation. **A.** Graph shows the expression of Piezo1 and 2 quantifed by RNAseq. **B.** Graph shows the relative expression of Piezo1 quantified by RT-qPCR in siRNA transfected cells (Unpaired t-test, siCTL – siPiezo1: p value 0.0164.) **C.** Schematic representation of the experimental design. Graph shows percentage of CTC extravasation in siRNAs-transfected cells at 21 hpi (Kruskal-Wallis test multiple comparison to siCtrl, siPiezo1: >0,9999; siPiezo2: 0,1314; siCtrl x2 >0,9999; siPiezo1+2: 0,2664; siCtrl: 34 embryos; siPiezo1: 48 embryos; siPiezo2: 34 embryos; siCtrl x2: 35 embryos; siPiezo1+2: 34 embryos). **D.** Confocal z-stack projections displaying representative examples of siCtrl and siPiezo-transfected cells at 24hpi in embryos. EC channel is displayed using inverted LUT to facilitate visualization. Zoom boxes show a single confocal plane to improve visualization of intravascular (yellow asterisks) and extravascular (red arrowhead) CTCs.

**Movie 1:** *EC-actomyosin dynamics mediate CTC extravasation*. Time-lapse z-stack projections show a CTC extravasating from an ISV. EC actin in black, EC nuclei in red, and CTC in magenta. Later in the movie, individual actin channel is displayed to facilitate the visualization of actin clustering.

**Movie 2:** Time-lapse confocal section showing EC calcium firing (signal displayed using fire LUT to facilitate the visualization of signal intensity, where blue is the minimum and white is the maximum) in a non-injected control embryo.

**Movie 3:** Time-lapse confocal section showing EC calcium firing (signal displayed using fire LUT to facilitate the visualization of signal intensity, where blue is the minimum and white is the maximum) and CTCs (white).

